# Parkinson’s Disease-linked LRRK2 structure and model for microtubule interaction

**DOI:** 10.1101/2020.01.06.895367

**Authors:** Colin K. Deniston, John Salogiannis, Sebastian Mathea, David M. Snead, Indarjit Lahiri, Oscar Donosa, Reika Watanabe, Jan Böhning, Andrew K. Shiau, Stefan Knapp, Elizabeth Villa, Samara L. Reck-Peterson, Andres E. Leschziner

## Abstract

**L**eucine **R**ich **R**epeat **K**inase **2** (*LRRK2*) is the most commonly mutated gene in familial Parkinson’s disease. LRRK2 is proposed to function in membrane trafficking and co-localizes with microtubules. We report the 3.5Å structure of the catalytic half of LRRK2, and an atomic model of microtubule-associated LRRK2 built using a reported 14Å cryo-electron tomography *in situ* structure. We propose that the conformation of LRRK2’s kinase domain regulates its microtubule interaction, with a closed conformation favoring binding. We show that the catalytic half of LRRK2 is sufficient for microtubule binding and blocks the motility of the microtubule-based motors kinesin and dynein *in vitro*. Kinase inhibitors that stabilize an open conformation relieve this interference and reduce LRRK2 filament formation in cells, while those that stabilize a closed conformation do not. Our findings suggest that LRRK2 is a roadblock for microtubule-based motors and have implications for the design of therapeutic LRRK2 kinase inhibitors.

## Introduction

Mutations in Leucine-Rich Repeat Kinase 2 (*LRRK2*) are the most common cause of familial Parkinson’s Disease (PD)^1^. LRRK2 is also linked to the idiopathic form of PD: mutations in *LRRK2* are a genetic risk factor^1^ and increased LRRK2 kinase activity is linked to disease^2^. A growing body of evidence suggests that mutations in LRRK2 are also risk factors for Crohn’s Disease and leprosy^3–5^. Like many other PD-causing genes, *LRRK2* is implicated in membrane trafficking^6^. Mutant LRRK2 causes defects in the trafficking of endosomes, lysosomes, autophagosomes, and mitochondria^6,7^. Furthermore, Rab GTPases, central regulators of membrane trafficking, are physiological substrates of LRRK2^8,9^. In cells, all five major PD-causing mutations increase the phosphorylation of LRRK2’s Rab substrates^8,9^. Long-distance transport of Rab-marked membranes occurs along microtubules and is driven by the microtubule-based motors dynein and kinesin^10,11^. LRRK2 function and/or pathology has also been linked to microtubules as LRRK2 partially co-localizes with microtubules in cells^12^ and four of the five major PD-causing mutations (Fig. 1a)^1^ enhance microtubule association of LRRK2^13^.

**Figure 1.**
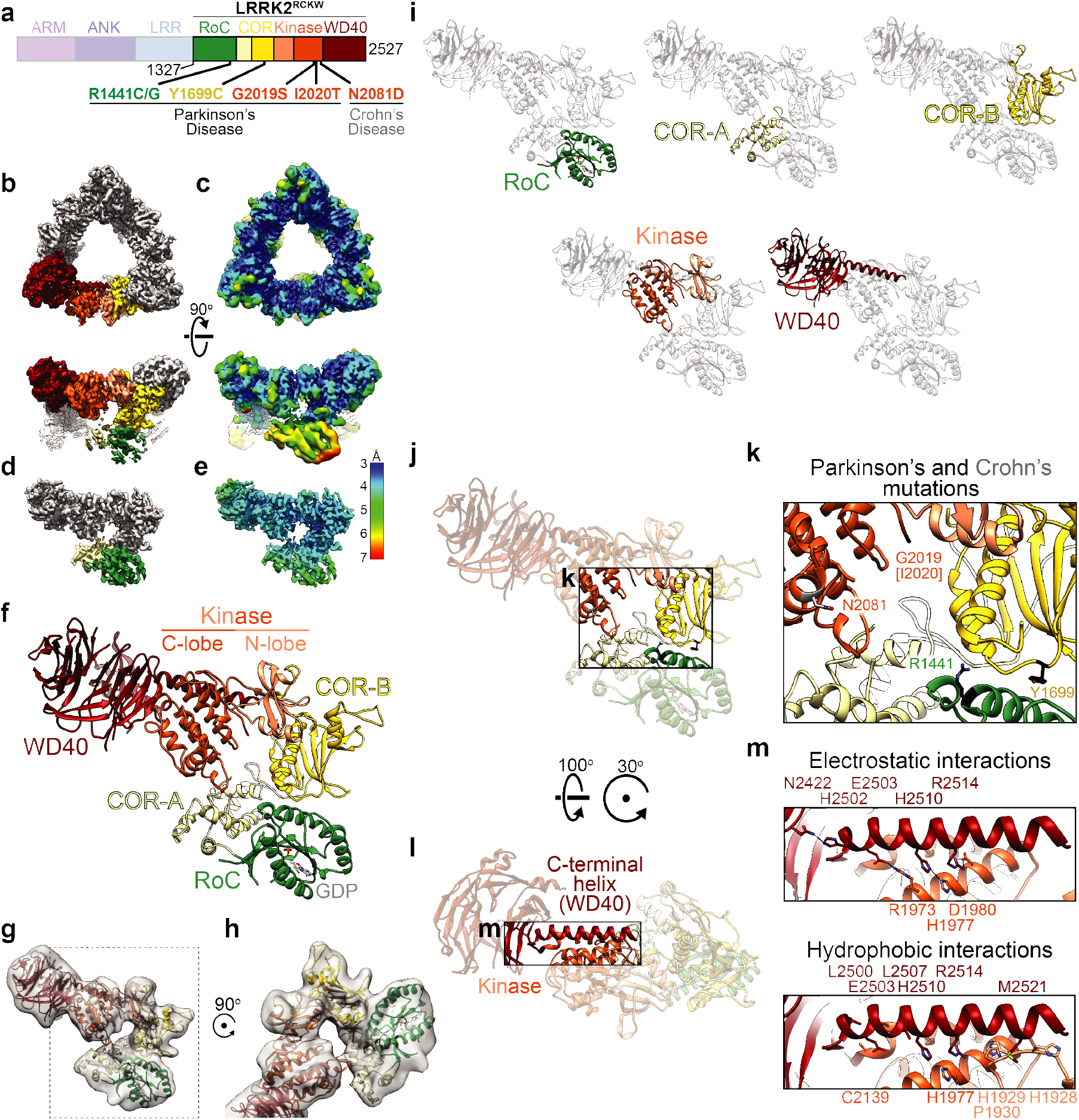
Cryo-EM structure of LRRK2^RCKW^. **a**, Schematic of the construct used in this study, with amino acid numbers of the beginning and end indicated. The N-terminal half of LRRK2, absent from our construct, is shown in dim colors. The same color-coding of domains is used throughout the paper. The five major familial Parkinson’s Disease mutations and a Crohn’s Disease-linked mutation are indicated below the diagram. **b**, 3.5Å map of the LRRK2^RCKW^ trimer, with one monomer highlighted using the colors shown in (a). **c**, Local resolution of the map shown in (b). **d**, Density for a LRRK2^RCKW^ monomer obtained after processing data where the RoC and COR-A domains were signal-subtracted from two of the monomers in each trimer (see Methods and Extended Data Fig. 2 for details). **e**, Local resolution for the monomer in (d), showing the improvements in the RoC and COR-A domains. **f**, Ribbon diagram of the atomic model of LRRK2^RCKW^. **g**, An 8.1Å cryo-EM map of monomeric LRRK2^RCKW^ with the atomic model built from the trimer docked in. **h**, Close up of the area highlighted by the dashed box in (g). **i**, Each LRRK2^RCKW^ domain is highlighted in color in the context of an otherwise grey ribbon representation of the full structure. **j**, **k**, Location of the Parkinson’s and Crohn’s Disease mutations listed in (a). **l**, Interface between the C-terminal helix and the kinase domain in LRRK2^RCKW^. **m**, details of the electrostatic and hydrophobic interactions found at that interface, with the residues involved indicated.

LRRK2 is a large (288 kDa) multi-domain protein. Its aminoterminal half is comprised of repetitive protein interaction motifs (Armadillo, Ankyrin, and Leucine-Rich Repeats) and its carboxyterminal catalytic half contains a Ras-like GTPase (Ras-of-Complex, or RoC domain), a kinase domain, and two other domains (C-terminal Of Roc, or COR, and WD40) (Fig. 1a). Despite LRRK2’s fundamental importance for understanding and treating PD, there is limited structural information on LRRK2. While structures are available for bacterial homologs of the LRR, RoC and COR domains^14,15^, and a *D. discoideum* homolog of the kinase domain^16^, the only two high-resolution structures available for the human protein are for its RoC^17^ and WD40 domains^18^ and no larger structures have been reported. Structures of the full-length protein obtained using negative stain^19^ and cryo-electron microscopy (Cryo-EM)^20^ have been published, but their resolutions were limited (22Å and 16Å, respectively). A recent reconstruction of LRRK2 bound to microtubules in cells using cryo-electron tomography (Cryo-ET) and sub-tomogram averaging led to a 14Å structure and proposed model of the catalytic half of LRRK2^21^. Here, we set out to determine a high-resolution structure of LRRK2’s catalytic half using cryo-EM, as well as to understand how it interacts with microtubules and how this impacts the movement of microtubule-based motors.

## Results

### Cryo-EM structure of the catalytic half of LRRK2

High-resolution studies on human LRRK2 have been limited by the lack of efficient expression systems resulting in stable LRRK2 protein. We tested a large number of constructs (Extended Data Fig. 1a), leading to the identification of one consisting of the carboxy-terminal half of LRRK2 (amino acids 1,327 to 2,527), which expressed well in insect cells and resulted in a stable protein (Extended Data Fig. 1b, c). This construct comprises the **R**oC, **C**OR, **k**inase and **W**D40 domains of LRRK2 (Fig. 1a), which we refer to as LRRK2^RCKW^. The COR domain was previously defined as consisting of two subdomains, COR-A and COR-B, based on its structure from a bacterial homolog^14^.

We determined a 3.5Å structure of LRRK2^RCKW^ in the presence of GDP using cryo-EM (Fig. 1b, c and Extended Data Fig. 2). On our grids, we observed a mixture of monomers, dimers and head-to-tail trimers and we used the trimer to solve the structure (Fig. 1b and Extended Data Fig. 2b). This trimer species was critical for reaching high resolution, but it is likely specific to the EM grid preparation as LRRK2^RCKW^ is predominantly monomeric, with a smaller percentage of dimers, in solution (Extended Data Fig. 3). The RoC and COR-A domains were flexible in our structure, yielding significantly lower resolutions than the rest of the protein as a result of the symmetry imposed on the trimer (Fig. 1b, c). In order to improve the density of this part of the map, we used signal subtraction to generate LRRK2^RCKW^ trimers containing only one of the three RoC and COR-A domains and subjected those to 3D classification and refinement focused on the monomer containing RoC and COR-A. This resulted in a 3.8Å structure with improved local resolution for RoC and COR-A (Fig. 1d, e, Extended Data Fig. 2f-i). We used a combination of Rosetta^22^ and manual building in coot^23^ to generate the final model. The RoC and COR-A domains were built using the signal subtracted maps, and then combined with the COR-B, kinase and WD40 domains, which were built using the original higher resolution trimer map (Fig. 1f and Extended Data Video 1). Importantly, the atomic model of LRRK2^RCKW^ we obtained from the trimers fits well into an 8.1Å reconstruction of a LRRK2^RCKW^ monomer (Fig. 1g, h), indicating that formation of the trimer does not cause major structural changes in the protein.

LRRK2^RCKW^ adopts an overall J-shaped structure, with the WD40, kinase and COR-B domains arranged along one axis, and CORA and RoC turning around back towards the kinase, bringing COR-A, and therefore the tightly associated RoC domain, which has GDP bound, in close proximity to the kinase’s C-lobe (Fig. 1f, i and Extended Data Video 1). This arrangement likely underpins the reported crosstalk between LRRK2’s kinase and GTPase^24,25^ (reviewed in^26^). Part of the FERM domain in the FAK-FERM complex approaches the FAK C-lobe in a similar way^27^ (Extended Data Fig. 4a, b). The RoC, COR-A and COR-B domains are arranged as previously seen in crystal structures of the COR^14^, RoC-COR^14,28^, and LRR-RoC-COR^15^ fragments of bacterial homologs of LRRK2. The N-lobe of the LRRK2^RCKW^ kinase domain, in particular its aC helix, forms an extensive interaction with the COR-B domain, with COR-B occupying a location reminiscent of that of Cyclin A in CDK2-Cyclin A^29^ (Extended Data Fig. 4a, c).

The kinase in LRRK2^RCKW^ is in an open, inactive conformation. Its activation loop is disordered beyond G2019, the location of one of the major familial PD mutations (G2019S) (Fig. 1j, k and Extended Data Video 2). Thus, I2020, the location of the other familial PD mutation found in the activation loop (I2020T), is disordered in our structure (Fig. 1j, k and Extended Data Video 2). R1441 and Y1699 are the sites of three other familial PD mutations and are located at the RoC-COR-B interface (Fig. 1j, k and Extended Data Video 2). Y2018 of the DYG motif (DFG motif in other kinases) is within hydrogen-bonding distance of the backbone carbonyl of I1933 in the N-lobe (Extended Data Fig. 4j, k). Given that a Y2018F mutation leads to hyperactivation of LRRK2’s kinase^30^, it is possible that a Y2018-I1933 hydrogen bond keeps the activation loop in an inactive conformation.

A unique feature of LRRK2 is a 28-amino acid α-helix located at its extreme C-terminus, following the WD40 domain (Fig. 1i, l, m and Extended Data Video 3). This helix extends along the entire kinase domain, interacting with both its C- and N-lobes (Fig. 1l, m and Extended Data Video 3). Deletion of this helix resulted in an insoluble protein (Extended Data Fig.1). While a number of other kinases have alpha helices in the same general location as LRRK2’s C-terminal helix, none of those interactions are as extensive as that observed in LRRK2 (Extended Data Fig. 4d-i). A residue near its end (T2524) is a known phosphorylation site for LRRK2^31^. Given the close proximity between T2524 and the N-lobe of the kinase domain, as well as the adjacent COR-B domain, we hypothesize that phosphorylation of this residue may play a role in regulating the kinase. Since the last two residues of the C-terminal helix are disordered in our structure, as is a neighboring loop in COR-B, it is possible that conditions exist where these regions become ordered and turn the C-terminal helix into an anchoring element that connects COR-B, the kinase and the WD40 domain.

Although our construct lacks the N-terminal half of full-length LRRK2, we were able to model the Leucine-Rich Repeats (LRR) by using a recent crystal structure of the LRR, RoC, and COR domains of *C. tepidum’s* Roco protein, a bacterial homolog of LRRK2^15^ (Extended Data Fig. 5a-e). Aligning the RoC and COR domains of this structure to those of LRRK2^RCKW^ resulted in a good fit for the LRR, which in the model wraps around the N-lobe of the kinase and comes in close proximity of its C-lobe as it connects to the RoC domain (Extended Data Fig. 5a-e). This model places the known S1292 autophosphorylation site in the LRR in close proximity to the kinase’s active site, and the Crohn’s disease-related residue N2081, located in the kinase’s C-lobe adjacent to the LRR (Extended Data Fig. 5f), suggesting the functional relevance of this potential interface.

### An atomic model of microtubule-bound LRRK2 filaments

Recently, a 14Å structure of microtubule-associated filaments of fulllength LRRK2 (carrying the filament-promoting I2020T mutation^13^) was reported using *in situ* cryo-ET and sub-tomogram averaging^21^ (Fig. 2a). The LRRK2 filaments formed on microtubules were right-handed^21^. The fact that microtubules are left-handed, combined with the absence of strong density connecting the LRRK2 filament to the microtubule surface^21^ raised the question of whether LRRK2’s interaction with microtubules is direct or is mediated by other microtubule-associated proteins. To address this, we combined purified microtubules with purified LRRK2^RCKW^ and observed them by cryo-EM. The resulting helical microtubule decoration (Fig. 2b) suggests that the interaction between LRRK2 and microtubules is direct and that the catalytic C-terminal half of LRRK2 is sufficient for the formation of microtubule-associated filaments.

**Figure 2.**
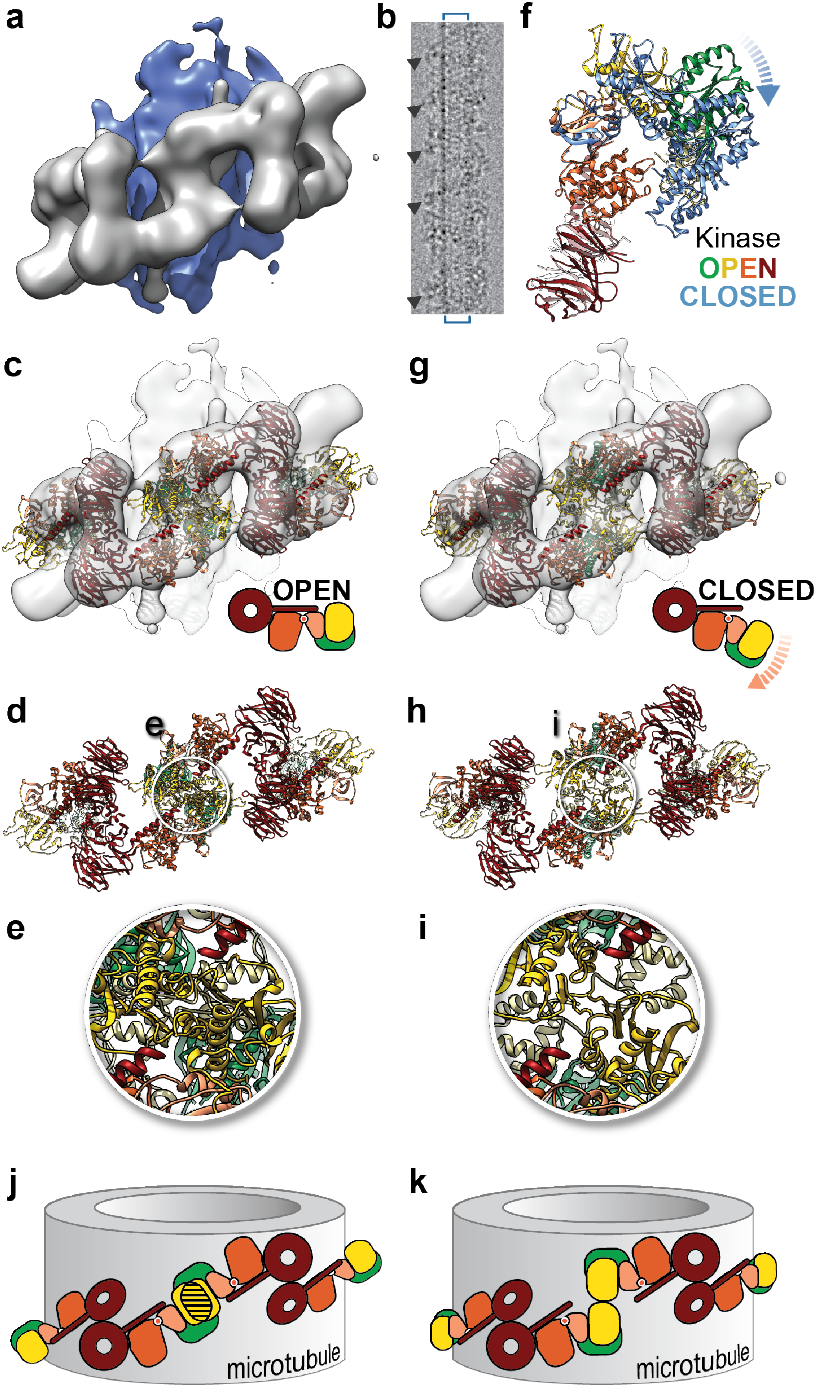
Modeling the microtubule-associated LRRK2 filaments. **a**, 14Å sub-tomogram average of a segment of microtubule-associated LRRK2 filament in cells. The microtubule is shown in blue and the LRRK2 filament in grey. **b**, Cryo-EM of microtubule-associated LRRK2^RCKW^ filaments reconstituted *in vitro* using purified LRRK2^RCKW^ and purified microtubules. The blue square brackets indicate the boundaries of the microtubule, and the arrowheads point to examples of LRRK2^RCKW^ filaments. **c**, Fitting of the LRRK2^RCKW^ structure, which has its kinase in an open conformation, into the sub-tomogram average. **d**, Atomic model of the LRRK2^RCKW^ filaments (c) with the sub-tomogram average density removed. The white circles highlight the filament interface mediated by interactions between COR domains, where clashes are found. **e**, Magnified views of the circled area shown in (d). **f**, Superposition of the LRRK2^RCKW^ structure (colored by domains) and a model of LRRK2^RCKW^ with its kinase in a closed conformation in blue (see Methods and Extended Data Fig. 6f-j for details on the model building). The dashed blue arrow indicates the general direction of movement upon closing of the kinase. **g**, Fitting of the closed-kinase model of LRRK2^RCKW^ into the subtomogram average. **h**, Atomic model of the closed-kinase LRRK2^RCKW^ filaments (g) with the sub-tomogram average density removed and a white circle highlighting the same interface as in (d). **i**, Magnified views of the circled area shown in (h). **j, k,** Cartoon representation of the two filament models, highlighting the clashes observed with open-kinase LRRK2^RCKW^ (j) and resolved with the closed-kinase model (k). Backbone clashes in (d, e) and (h, i) were measured in Chimera with Find Clashes using a polyalanine model of LRRK2^RCKW^. There were 997 clashes in the filament modeled using our LRRK2^RCKW^ structure (“open” form (d, e)) and 184 clashes in the filament built using the closed-kinase model of LRRK2^RCKW^ (h, i), corresponding to an overall reduction of 81.8% in the number of clashes.

Previously, integrative modeling was used to build a model into the *in situ* structure of microtubule-associated LRRK2^21^. This modeling indicated that the observed density was comprised of the RoC, COR, Kinase and WD40 domains and gave orientation ensembles for each domain^21^ that were in good agreement with the high-resolution structure of LRRK2^RCKW^ presented here. However, given the uncertainties in the domain orientations intrinsic to the integrative modeling approach, we set out to build an atomic model of the microtubule-bound LRRK2 filaments by combining our 3.5Å structure of LRRK2^RCKW^ with the 14Å *in situ* structure of microtubule-associated LRRK2. To guide model building, we took advantage of the characteristic shape of the WD40 domains, which are a prominent feature of the *in situ* structure (Fig. 2a and Extended Data Fig. 6a-c) and that a crystal structure of a WD40 dimer from LRRK2 was recently reported^18^. We first replaced the WD40 domains in the crystal structure with those from our LRRK2^RCKW^ structure (there are small differences between them) (Extended Data Fig. 6a) and then docked this WD40 dimer into the sub-tomogram average density (Extended Data Fig. 6b). Next, we aligned the LRRK2^RCKW^ structure to it, thus imposing the WD40 dimer interface from the crystal structure on our model (Extended Data Fig. 6d, e). This initial model revealed that the LRRK2^RCKW^ structure is sufficient to account for the density seen in the *in situ* structure (Fig. 2a, c), in agreement with our ability to reconstitute microtubule-associated LRRK2^RCKW^ filaments *in vitro* (Fig. 2b), and with the earlier integrative modeling^21^.

Although the LRRK2^RCKW^ structure fits the overall shape of the sub-tomogram average, we noticed significant clashes at the filament interface formed by the COR domains (Fig. 2d, e). Since the kinase in our LRRK2^RCKW^ structure is in an open conformation (Fig. 2c), we wondered whether filament formation might require LRRK2’s kinase to be in a closed conformation. We modeled a kinase-closed LRRK2^RCKW^ by running structural searches (DALI server) with either the N- or C-lobes of LRRK2^RCKW^’s kinase and looking for a kinase in a closed conformation whose N- and C-lobes best matched those of LRRK2^RCKW^. The top candidate was Interleukin-2 inducible T-cell kinase (ITK) (PDB: 3QGY). We aligned ITK to LRRK2^RCKW^ using the C-lobes of the two kinases, and then aligned the N-lobe of LRRK2^RCKW^’s kinase to that of ITK, moving RoC, COR-A and COR-B as a rigid body along with the kinase’s N-lobe (Fig. 2f and Extended Data Fig. 6f-j). We then repeated the docking into the *in situ* structure, this time using the model of kinase-closed LRRK2^RCKW^. In addition to improving the fit visually (Fig. 2g, h), using the kinase-closed LRRK2^RCKW^ model resolved more than 80% of the backbone clashes we had observed with our kinase-open LRRK2^RCKW^ structure (Fig. 2i). A closed conformation for the kinase had already been proposed by the earlier integrative modeling^21^. Given this data, we hypothesize that the conformation of LRRK2, as driven by the kinase, controls its ability to associate with microtubules, with a closed kinase promoting oligomerization, which increases binding, and an open (inactive) one disfavoring it (Fig. 2j, k).

The LRRK2 filaments in our kinase-closed model are formed by two types of homotypic interactions, each resulting in a two-fold axis of symmetry perpendicular to the microtubule axis: one is mediated by the WD40 domain, and the other by the COR-A and COR-B domains (Fig. 3a-d). Similar interfaces were reported based on integrative modeling performed with the 14Å LRRK2 structure obtained by cryo-ET^21^. We wondered whether these interfaces were specific to the microtubule-associated form of LRRK2. Additional structures we obtained from our cryo-EM data showed this was not the case; in addition to the trimers we used to obtain the 3.5Å structure of LRRK2^RCKW^, grids prepared in the absence of microtubules also contained dimers that were mediated by the same two interfaces seen in the filaments (Fig. 3e, f and Extended Data Figs. 7-10). We observed these dimers under different conditions, both in the presence and absence of kinase inhibitors (Fig. 3e, f and Extended Data Fig. 10). All of our cryo-EM maps of dimers fit well the molecular models of dimers derived from the filaments, suggesting that the same interfaces observed off microtubules are involved in the formation of the microtubule-associated filaments.

**Figure 3.**
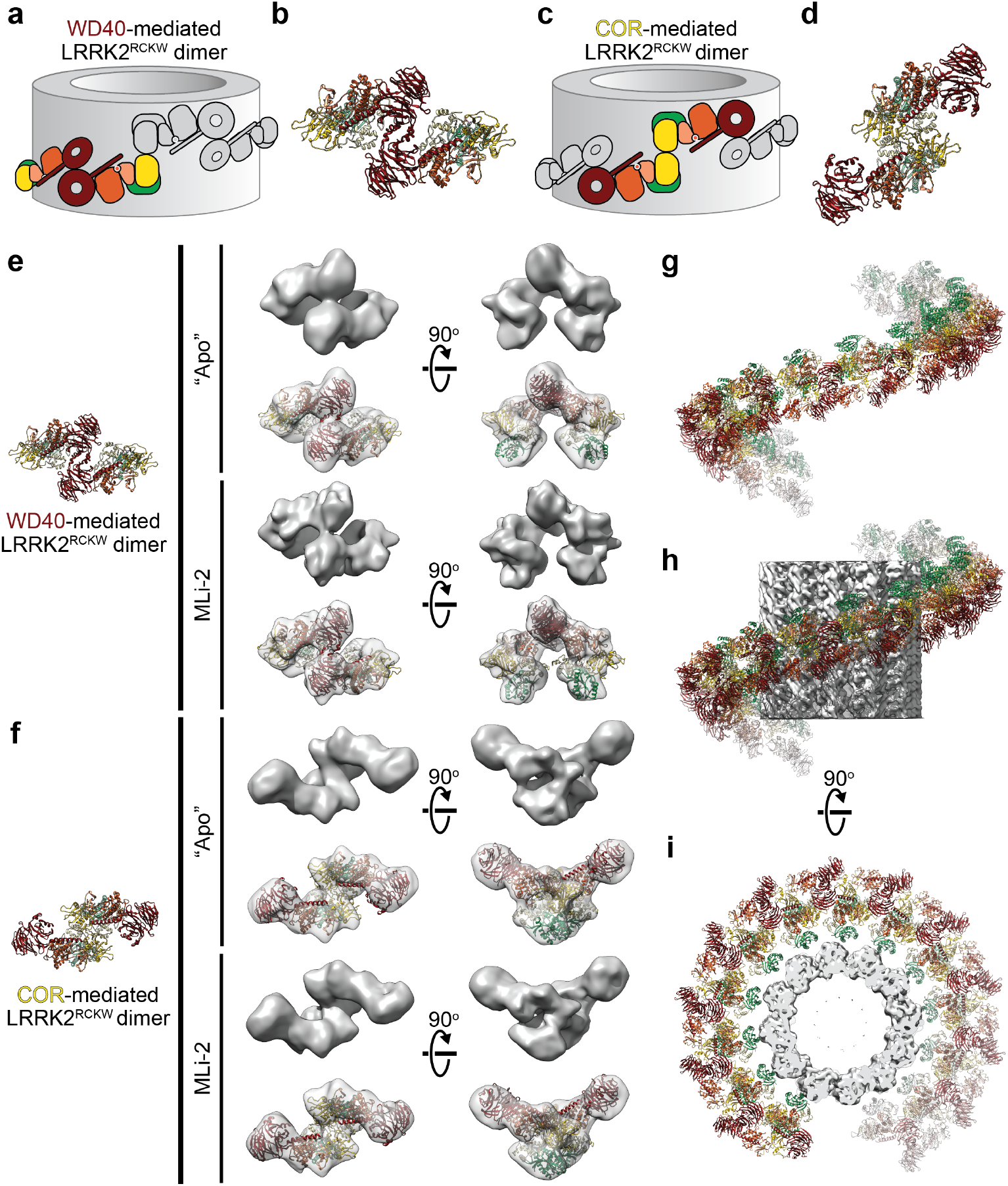
LRRK2^RCKW^ forms WD40- and COR-mediated dimers outside the filaments. **a-d**, The same filament model shown in Fig. 2j, k is shown here in grey, with either a WD40-mediated (a), or COR-mediated (c) LRRK2^RCKW^ dimer highlighted with domain colors. The corresponding molecular models are shown next to the cartoons (b, d). **e, f,** Cryo-EM reconstructions of LRRK2^RCKW^ dimers obtained in the absence of inhibitor (“Apo”), or in the presence of MLi-2. The molecular models (left) are the same ones shown in (b, d) for the WD40-mediated (e) and COR-mediated (f) LRRK2^RCKW^ dimers. The models are shown in an orientation that matches the cryo-EM maps shown in the first column. For each reconstruction, two orientations of the map are shown: down the two-fold axis at the dimerization interface (left), and perpendicular to it (right). For each species, the top row shows the cryo-EM map and the bottom row a transparent version of the map with a model docked in. The models docked into the maps obtained in the presence of MLi-2 are those derived from building the filaments (Fig. 2g-i). Those docked into the maps obtained in the absence of inhibitor were built with the LRRK2^RCKW^ structure (*i.e*. with an open kinase) but maintaining the interfaces identified in the filaments. Both types of dimers were also observed in the presence of Ponatinib but the preferred orientation of the sample prevented us from obtaining a 3D reconstruction; Extended Data Fig. 10 shows 2D class averages instead. **g,** We built molecular models of the WD40-mediated and COR-mediated dimers of LRRK2^RCKW^ obtained in the presence of MLi-2 (e, f) by fitting the two halves of LRRK2^RCKW^ split at the junction between the N- and C-lobes of the kinase (see Extended Data Fig. 11). We then aligned, in alternating order, copies of these dimers. This panel shows the resulting right-handed helix. **h, i,** The helix has dimensions compatible with the diameter of a 12-protofilament microtubule (EMD-5192)^53^, which was the species used to obtain the tomographic reconstruction shown in Fig. 2a^21^, and has its RoC domains pointing towards the microtubule surface.

Separately, we docked LRRK2^RCKW^, split in half at the junction between the N- and C-lobes of the kinase, into the cryo-EM map of a WD40-mediated dimer obtained in the presence of MLi-2 (Extended Data Fig. 11a-c). We docked the resulting closed-kinase model of LRRK2^RCKW^ (Extended Data Fig. 11e) into the cryo-EM maps of WD40- and COR-mediated dimers obtained in the presence of MLi-2 to generate molecular models of these dimers (Extended Data Fig. 11c, d-g). Finally, we aligned these models, in alternating order, to build a polymer *in silico*. The resulting structure was a right-handed helix with the same general geometric properties seen in the cellular LRRK2 filaments, indicating that those properties are largely encoded in the structure of LRRK2^RCKW^ itself (Fig. 3g-i and Extended Data Fig. 11g, h). Docking the same two halves of LRRK2^RCKW^ into the cryo-EM map of a monomer obtained in the absence of inhibitors led to a structure very similar to that obtained from the trimers, further confirming that formation of the trimer did not alter the conformation of LRRK2^RCKW^ (Extended Data Fig. 11a, b, d, e).

These data, along with the apparent lack of any residue-specific interactions between LRRK2 and the microtubule, suggest that the microtubule may be providing a surface for LRRK2 to oligomerize using interfaces that exist in solution. Consistent with this idea, the surface charge of the microtubule facing the LRRK2^RCKW^ filament is acidic, while there are a number of basic patches on the LRRK2^RCKW^ filament facing the microtubule (Extended Data Fig. 12). The unstructured C-terminal tails of α- and β-tubulin, which were not included in the surface charge calculations, are also acidic. Finally, the interface in the COR-mediated dimer we observed for LRRK2^RCKW^ differs from that reported for the homologous Roco protein from *C. tepidum^14,15^* (Extended Data Fig. 13). While the GTPase domains interact directly in the dimer of the bacterial protein^15^, they are not involved in the dimerization interface of LRRK2 (Extended Data Fig. 13d, e).

### LRRK2^RCKW^ inhibits kinesin and dynein motility in vitro

To test our hypothesis that the conformation of LRRK2’s kinase domain regulates its interaction with microtubules, we needed a sensitive assay to measure the association of LRRK2^RCKW^ with microtubules and a means to control the conformation of its kinase. Microtubule association was monitored by measuring the effect of LRRK2^RCKW^ on the movement of microtubule-based motors. We used a truncated dimeric human kinesin-1, Kif5B (“kinesin” here)^32^, which moves towards the plus ends of microtubules, and activated human cytoplasmic dynein-1/ dynactin/ninein-like complexes (“dynein” here)^33^, which move in the opposite direction. Using single-molecule *in vitro* motility assays (Fig. 4a) we found that low nanomolar concentrations of LRRK2^RCKW^ inhibited both kinesin and dynein movement, with near complete inhibition achieved at 25 nM LRRK2^RCKW^ (Fig. 4b-e). We hypothesized that LRRK2^RCKW^ was acting as a roadblock for the motors. In agreement with this hypothesis, the run length of kinesin was reduced (Fig. 4f), while its velocity remained relatively constant (Fig. 4g). Activated cytoplasmic dynein-1 complexes also showed a significant reduction in run length in the presence of LRRK2^RCKW^ (Fig. 4h). Thus, LRRK2 ^RCKW^ robustly blocked the motility of both kinesin and dynein motors *in vitro*.

**Figure 4.**
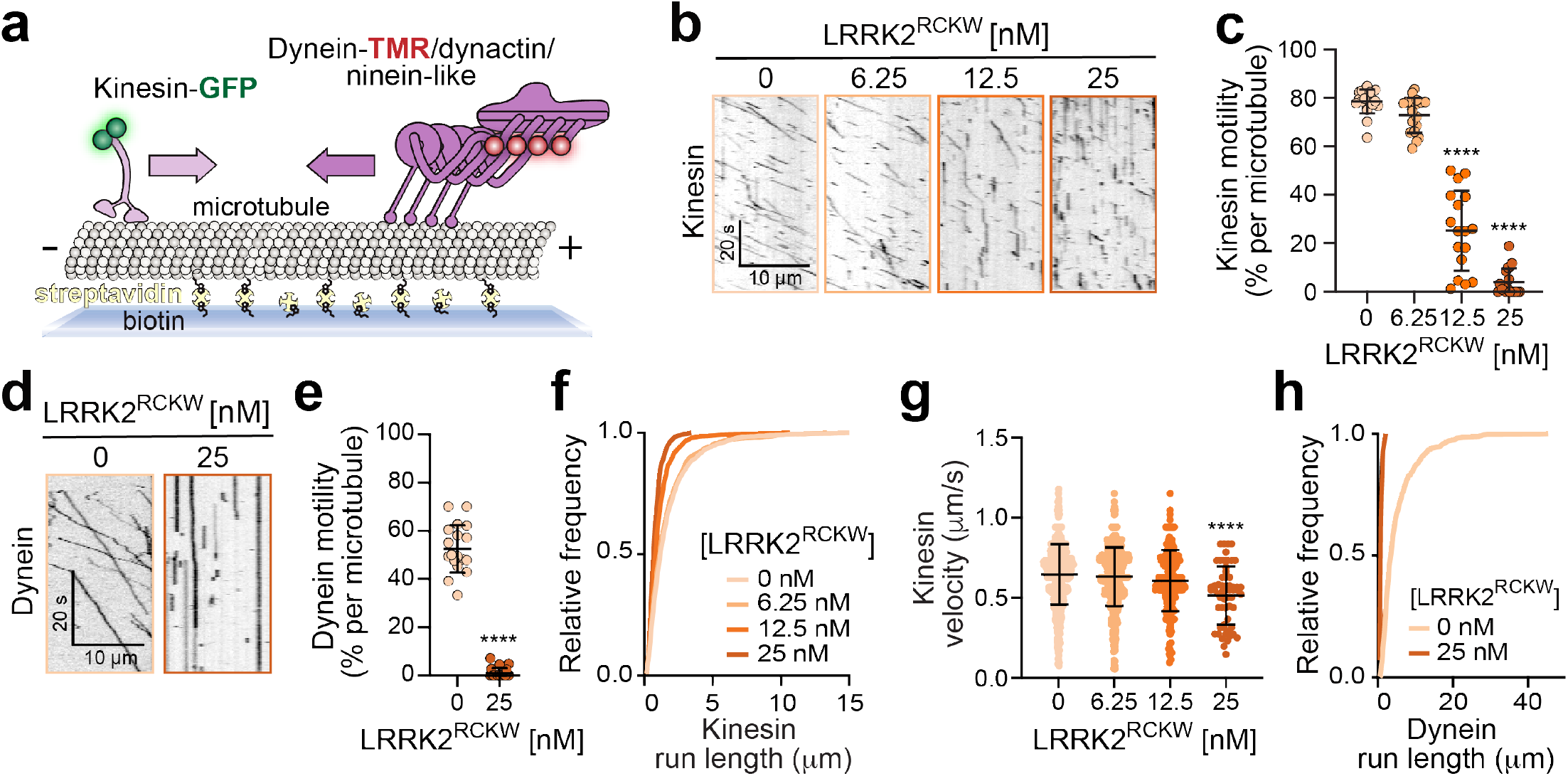
LRRK2^RCKW^ inhibits the motility of the microtubule-based motors kinesin-1 and cytoplasmic dynein-1. **a**, Schematic representation of the experimental setup for the single-molecule motility assays. See Methods for details. **b**, Example kymographs showing that increasing concentrations of LRRK2^RCKW^ reduce kinesin-1-GFP (kinesin) runs. **c**, The percentage of motile kinesin events per microtubule as a function of LRRK2^RCKW^ concentration. Data are mean ± s.d. (n = 20, 18, 17, and 18 microtubules from left to right; quantified from two independent experiments). ****p < 0.0001 calculated using the Kruskal-Wallis test with Dunn’s posthoc for multiple comparisons. **d,** Example kymographs showing that 25 nM LRRK2^RCKW^ reduces dynein/ dynactin/ ninein-like (dynein) runs. **e,** The percentage of motile dynein events per microtubule as a function of LRRK2^RCKW^ concentration. Data are mean ± s.d. (n = 19 and 21 microtubules from left to right; quantified from two independent experiments). ****p < 0.0001 calculated using the Mann Whitney test. **f,** Cumulative frequency distribution of run lengths of kinesin as a function of LRRK2^RCKW^ concentration. From top to bottom: n = 1166, 1090, 571, and 213 runs. Mean decay constants (tau) ± 95% confidence intervals; microns are 1.667 ± 0.05, 1.570 ± 0.046, 1.048 ± 0.088, and 0.813 ± 0.145. Data quantified from two independent experiments. **g**, Velocity of kinesin as a function of LRRK2^RCKW^ concentration. Data are mean ± s.d. (from left to right: n = 680, 604, 228, and 59 runs quantified from two independent experiments). ****p < 0.0001 calculated using a one-way ANOVA with Dunn’s posthoc for multiple comparisons. All other conditions are *n.s*. **h,** Cumulative frequency distribution of run lengths for dynein as a function of LRRK2^RCKW^ concentration. (n = 659 and 28 runs from top to bottom; mean decay constants (tau) ± 95% confidence intervals; microns are 4.980 ± 0.147 and 0.8460 ± 0.415). Data quantified from two independent experiments. See Extended Data Table 1 for all source data and replicate information.

### Ponatinib and GZD-824 rescue motor inhibition by LRRK2^RCKW^ in vitro

Our hypothesis predicts that the closed conformation of LRRK2’s kinase domain will favor microtubule binding. Conversely, it predicts that conditions that stabilize the kinase in an open conformation will lead to decreased microtubule binding of LRRK2^RCKW^, resulting in relief of LRRK2^RCKW^-dependent inhibition of kinesin and dynein motility. To test these predictions, we searched for a Type 2 kinase inhibitor that binds tightly to LRRK2 with structural evidence that it stabilizes an open kinase conformation. We selected Ponatinib as our initial candidate inhibitor as it has a reported K_i_ for LRRK2 of 31 nM^34^, and crystal structures show it bound to RIPK2^35^ and IRAK4 in open conformations (Extended Data Fig. 14). We confirmed that Ponatinib also inhibited LRRK2^RCKW^ by monitoring phosphorylation of the known LRRK2 substrate, Rab8a^8^ *in vitro* (Extended Data Fig. 15a).

As our hypothesis predicted, Ponatinib rescued kinesin motility in a dose-dependent manner at concentrations of LRRK2^RCKW^ (25 nM) that had resulted in almost complete inhibition of the motors (Fig. 5a, Extended data Fig. 15b-e). We observed similar effects with GZD-824, a chemically-related Type 2 kinase inhibitor^36^ (Fig. 5a and Extended Data Fig. 15d, e). Our hypothesis also predicted that kinase inhibitors that stabilize the closed form of the kinase should be unable to rescue the motors and may even enhance the inhibitory effect of LRRK2^RCKW^ by increasing its interaction with microtubules. Indeed, the LRRK2-specific Type 1 inhibitors MLi-2^37,38^ and LRRK2-IN-1^39^, which are known^40^ or expected^37^ to stabilize a closed conformation of the kinase, further enhanced the inhibitory activity of LRRK2^RCKW^ on kinesin motility (Fig. 5a). As was the case with Ponatinib, GZD-824, MLi-2 and LRRK2-IN-1 all inhibited phosphorylation of Rab8a by LRRK2^RCKW^ (Extended Data Fig. 15a). Similar to kinesin, dynein motility was rescued by Ponatinib and GZD-824, but not MLi-2 or LRRK2-IN-1 (Fig. 5b and Extended Data Fig. 15d, f). These data suggest that the effects of LRRK2 kinase inhibitors on microtubule-based motility should be taken into account when designing LRRK2-targeted PD therapeutics.

**Figure 5.**
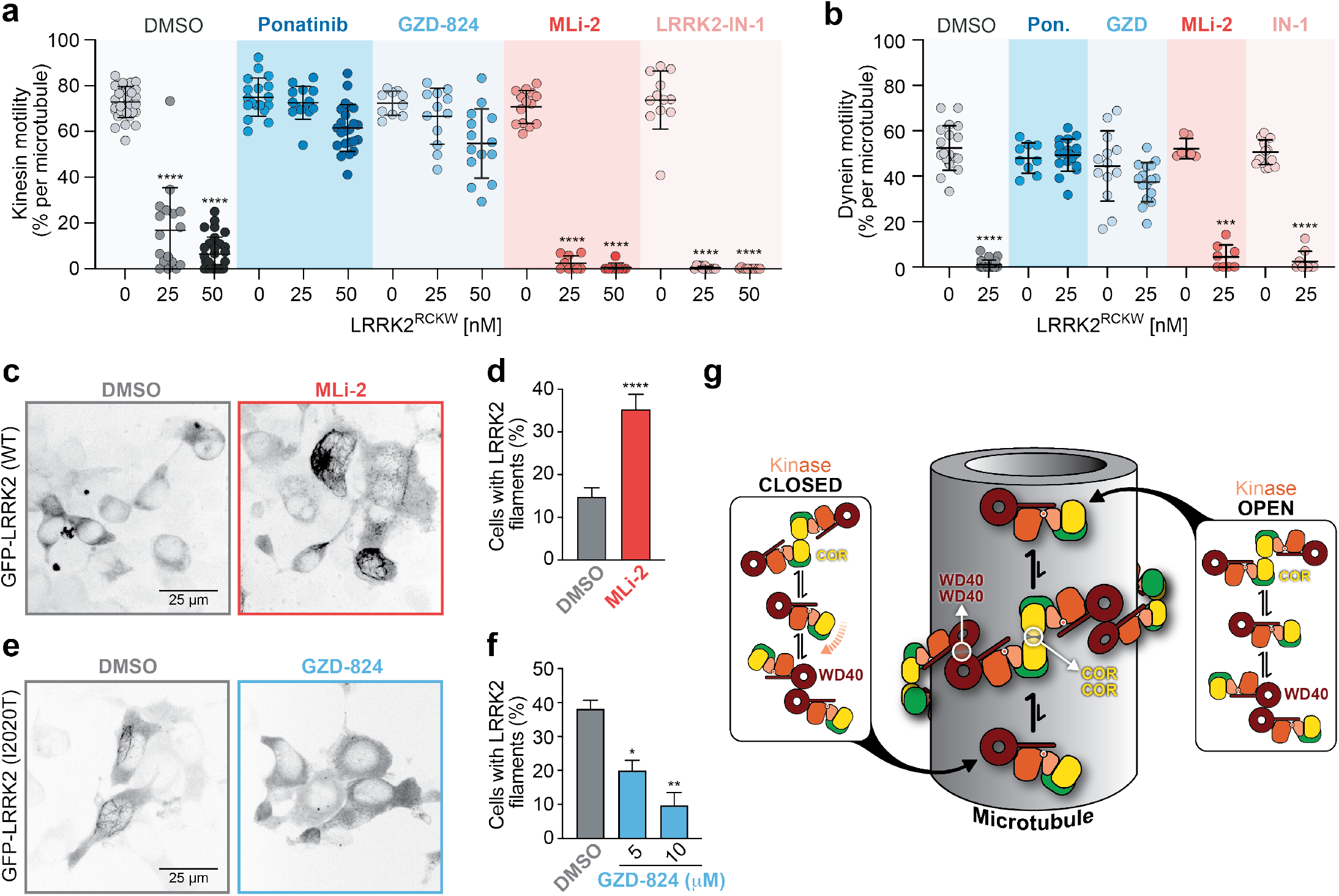
Type 2 kinase inhibitors, but not Type 1, rescue microtubule-based motor motility and reduce LRRK2 filament formation in cells. **a,** Effects of different kinase inhibitors on LRRK2^RCKW^’s inhibition of kinesin motility. Data is shown as the percentage of motile kinesin events per microtubule as a function of LRRK2^RCKW^ concentration in the absence of any kinase inhibitor (DMSO) or in the presence of the indicated kinase inhibitor (Ponatinib (Type 2): 10 μM; GZD-824 (Type 2): 10 μM; MLi-2 (Type 1): 1 μM; and LRRK2-IN-1 (Type 1): 1 μM). Data are mean ± s.d. (from left to right: n = 34, 17, 30, 18, 14, 23, 10, 12, 14, 14, 14, 8, 12, 11, and 11 microtubules quantified from two to four independent experiments). ****p < 0.0001 calculated using the Kruskal-Wallis test with Dunn’s posthoc for multiple comparisons (comparisons were within drug only). **b.** Same as (a) but with dynein/ dynactin/ ninein-like (dynein). DMSO conditions reproduced from Fig. 4c for comparison. (From left to right: n = 19, 21,9, 18, 13, 16, 7, 8, 14, and 9 microtubules quantified from two independent experiments). ***p < 0.001 and ****p < 0.0001 calculated using the Kruskal-Wallis test with Dunn’s posthoc for multiple comparisons (comparisons were within drug only). **c**, The Type 1 kinase inhibitor MLi-2 (500 nM) treated for 2 hrs increases WT GFP-LRRK2 filament formation in 293T cells compared to DMSO-treated control cells. **d,** Quantification of the experiment shown in (c). Data are mean ± s.e.m. (n = 10 [DMSO] and 6 for [MLi-2] technical replicates (~ 20 cells per replicate). ****p=0.0002 Mann Whitney test. **e,** The Type 2 kinase inhibitor GZD-824 (5 μM) treated for 30 mins decreases GFP-LRRK2 (I2020T) filament formation relative to DMSO-treated control cells. **f,** Quantification of the experiment shown in (e). Data are mean ± s.e.m. (n = 9 [DMSO], 8 [5 μM], and 4 [10 μM] technical replicates (> 40 cells per replicate). *p=0.0133 and **p=0.0012 calculated using Krustal-Wallis test with Dunn’s posthoc for multiple comparisons. See Extended Data Table 1 for all source data and replicate information. **g,** Schematic representation of our hypothesis that the conformation of LRRK2’s kinase controls its association with microtubules. LRRK2 (represented here by LRRK2^RCKW^) can have its kinase in either an open or closed conformation. The different species we observed (monomers and both COR- and WD40-mediated dimers) are represented in the rounded rectangles, but only monomers are shown on the microtubule for simplicity. Our model proposes that only the kinase-closed form of LRRK2 is compatible with the formation of microtubule-associated filaments. The figure shows a single turn of a LRRK2 filament to emphasize that we propose that shorter species will be the relevant ones under physiological conditions. The two interfaces that mediate filament formation, which are the same ones observed in the dimers, are indicated.

### GZD-824 reduces filament formation in cells

In cells, LRRK2 forms filaments that colocalize with a subset of microtubules and are sensitive to the microtubule depolymerizing drug nocodazole^13^. This association is enhanced by the PD-linked mutations R1441C, R1441G, Y1699C and I2020T^13,41^ and by Type 1 kinase inhibitors^39,42^. We tested our kinase conformation hypothesis in 293T cells by determining if Type 1 and Type 2 kinase inhibitors had opposite effects on the formation of cellular microtubule-associated LRRK2 filaments. Consistent with previous findings, the Type 1 inhibitor MLi-2 enhanced LRRK2’s-microtubule association (Fig. 5c, d), suggesting that the closed conformation of the kinase favors binding to microtubules in cells. In contrast, we found that the Type 2 inhibitor GZD-824 reduced the filament-forming ability of overexpressed LRRK2 (carrying the mutation I2020T; Fig. 5e, f). This reduction in LRRK2 filament formation was not due to changes in LRRK2 protein expression levels (Extended Data Fig. 16a, b) or the overall architecture of the microtubule cytoskeleton (Extended Data Fig. 16c).

## Discussion

Here we reported the 3.5Å structure of the catalytic half of LRRK2 (LRRK2^RCKW^), where most major PD-linked mutations are located. Our model represents the first structure of a large portion of human LRRK2 with sufficient resolution to yield molecular information about LRRK2’s activity and its regulation. LRRK2^RCKW^ is J-shaped, which brings its GTPase (RoC domain) and kinase in close proximity despite their separation along the linear sequence of LRRK2. The direct contact we observe between the C-lobe of the kinase and COR-A, which is closely associated with the GTPase, provides a structural context for the crosstalk between these two catalytic domains (reviewed in^26^). Our cryo-EM structure, obtained in the absence of kinase inhibitors, shows the kinase in an open (inactive) conformation. A unique feature of the structure is a long alpha helix located at the C-terminus of LRRK2, following the WD40 domain. Running along the long axis of the kinase domain, this helix has multiple interactions with both its N- and C-lobes. Given these interactions, and the presence of a known phosphorylation site near its end (T2524)^31^, it is likely that this helix plays an important role in regulating LRRK2’s kinase activity.

Our ability to generate microtubule-associated LRRK2^RCKW^ filaments using purified components demonstrates that these filaments can form in the absence of other proteins and that the C-terminal half of LRRK2 is sufficient for their assembly. This is also in agreement with the observation that our LRRK2^RCKW^ structure accounts for the filament density observed in a recent 14Å structure of LRRK2-microtubule associated filaments in cells obtained by cryo-ET^21^.

We used the high-resolution structure of LRRK2^RCKW^, in combination with the 14Å *in situ* structure^21^, to build an atomic model of the microtubule-associated LRRK2 filaments. Docking of our LRRK2^RCKW^ structure, with its open kinase domain, into the microtubule-associated *in situ* structure led to significant steric clashes. These clashes were largely resolved by modeling LRRK2’s kinase in a closed conformation, leading us to hypothesize that the conformation of LRRK2’s kinase controls its association with microtubules, with a closed conformation favoring its oligomerization on microtubules (Fig. 5g). In our model of microtubule-associated LRRK2^RCKW^ filaments, the filaments form through head-head and tail-tail interactions involving either the COR or WD40 domains of LRRK2. Formation of these interfaces does not require LRRK2 to interact with microtubules, as we obtained lower resolution cryo-EM structures of both types of LRRK2^RCKW^ dimers in the absence of microtubules. In addition, aligning atomic models of these dimers *in silico* resulted in a right-handed filament with similar geometric properties to the LRRK2 filaments observed in cells^21^. Thus, the ability of LRRK2 to form filaments is a property inherent to LRRK2, and specifically the RCKW domains. Since LRRK2^RCKW^ exists both as a monomer and dimer in solution, it remains to be determined what the minimal filament-forming unit is. We propose that the surface charge complementarity between the microtubule and LRRK2, as well as the size and shape of the microtubule stimulates the formation of LRRK2 filaments.

We tested the model that the conformation of LRRK2’s kinase regulates microtubule association both *in vitro* and in cells using kinase inhibitors that are known or expected to stabilize either the open (Type 2) or closed (Type 1) conformations of the kinase. In support of our model, Type 2 inhibitors relieved the LRRK2^RCKW^-dependent inhibition of the microtubule-based motors kinesin and dynein and reduced LRRK2 filament formation in cells, while Type 1 inhibitors failed to rescue the motors and enhanced filament formation in cells. In contrast to our structural studies, which used high concentrations of LRRK2^RCKW^ (cryo-EM) or overexpressed LRRK2 in cells (cryo-ET), our single-molecule motility assays showed that low nanomolar concentrations of LRRK2^RCKW^ negatively impact the microtubule-based motility of both kinesin and dynein *in vitro*. At these low concentrations it is likely that LRRK2 would not form the long, highly ordered filaments and microtubule bundles observed in cells overexpressing the protein; instead, we hypothesize that at endogenous expression levels in cells LRRK2 forms short stretches of filaments on microtubules.

Our data showing that increasing concentrations of LRRK2^RCKW^ dramatically shorten the distance kinesin moves (run length), without affecting its velocity suggest that LRRK2 acts as a roadblock. Other microtubule-associated proteins (MAPs), such as MAP2 and Tau, also inhibit kinesin motors^43^ (bioRxiv 10.1101/731604). In contrast to kinesin, dynein is largely unaffected by MAP2 and Tau^43^ (bioRxiv 10.1101/731604), likely due to its ability to sidestep to adjacent protofilaments^44–46^. LRRK2’s unusual ability to inhibit dynein motility may be a consequence of its forming oligomers that block dynein’s sidestepping.

What is the physiological role of non-pathogenic microtubule-associated LRRK2? Our data show that low nanomolar concentrations of LRRK2 act as a roadblock for microtubule-based motors. Dynein and kinesin have been shown to bind directly or indirectly to many Rab-marked cargos^47–51^. Our data also show that the microtubule-associated form of LRRK2 has its kinase in a closed (and potentially active) conformation. Given this, it is possible that when microtubule-associated LRRK2 stalls the movement of kinesin and dynein, this increases the likelihood that LRRK2 will phosphorylate cargo-associated Rab GTPases on their switch 2 region^8^, ultimately leading to effector dissociation^8^ and cargo release. This scenario suggests that increased microtubule binding by LRRK2 carrying PD mutations could (1) block membrane transport driven by molecular motors by acting as a roadblock, (2) lead to increased phosphorylation of physiological microtubule-associated substrates and/or phosphorylation of non-physiological substrates, or (3) a combination of the first two points. In support of this idea, the four familial PD mutations that enhance LRRK2 microtubule binding^13^ also show higher levels of Rab GTPase phosphorylation in cells than the G2019S LRRK2 mutant^8,9,52^, whose microtubule binding is not enhanced over wild-type LRRK2^13^. Testing this and other models is an important future direction to understand the cell biological function of both wild-type and pathogenic LRRK2.

Regardless of what role LRRK2’s binding to microtubules plays in PD, our data have important implications for the design of LRRK2 kinase inhibitors for therapeutic purposes. Our data predict that treatment with inhibitors that promote binding of LRRK2 to microtubules by favoring a closed conformation of its kinase will block microtubule-based trafficking, while inhibitors that favor an open conformation of the kinase will not.

## End Matter

### Author Contributions and Notes

CKD collected and processed the cryo-EM data. JS performed the singlemolecule and cellular assays with the help of OD. SM designed the LRRK2^RCKW^ construct and purified the protein. CKD and IL built the molecular model of LRRK2^RCKW^. DS performed the SEC-MALS and phosphorylation assays. RW and JB performed the cellular cryo-ET. AKS contributed to the structural analysis and provided advice on the selection of kinase inhibitors. SK, EV, SLR-P and AEL directed and supervised the research. CKD, JS, SLR-P and AEL wrote the manuscript and SM, DS, OD, AKS, SK and EV edited it.

The authors declare no conflict of interest.

Correspondence and material requests can be made to either Andres E. Leschziner or Samara L. Reck-Peterson.

## Acknowledgments

We thank Susan Taylor for her role in initiating this collaborative work, which was partially supported by multi-investigator grants from the Michael J Fox Foundation (grant #s: 11425 and 11425.02) (PI: Susan Taylor). We also thank the Nikon Imaging Center at UC San Diego, where the confocal microscopy was performed, the use of instruments at the Electron Imaging Center for NanoMachines supported by NIH (1S10RR23057, 1S10OD018111, and 1U24GM116792), NSF (DBI-1338135) and CNSI at UCLA, the UC San Diego Cryo-EM Facility, John P. Gillies and Agnieszka Kendrick for technical support with protein purifications, and Andrea Dickey for feedback on the manuscript. CKD was initially supported by the Molecular Biophysics Training Grant (NIH Grant T32 GM008326) and is currently supported by a Predoctoral Fellowship from the Visible Molecular Cell Consortium and Center for Trans-scale Structural Biology (UC San Diego). DS is supported by an A.P. Giannini Foundation postdoctoral fellowship. AKS receives salary and support from the Ludwig Institute for Cancer Research. EV is supported by a NIH Director’s New Innovator Award DP2GM123494. SLR-P is an investigator of the Howard Hughes Medical institute and is also supported by R01GM121772. AEL is supported by R01GM107214. SK is grateful for support from the SGC, a registered charity that receives funds from AbbVie, Bayer Pharma AG, Boehringer Ingelheim, Canada Foundation for Innovation, Eshelman Institute for Innovation, Genome Canada, Innovative Medicines Initiative [ULTRA-DD], Janssen, Merck KGaA Darmstadt Germany, MSD, Novartis Pharma AG, Ontario Ministry of Economic Development and Innovation, Pfizer, São Paulo Research Foundation-FAPESP, Takeda, and the Wellcome, as well as Boehringer Ingelheim for funding initial structural studies of this project. MS is supported by DFG grant HE1818/11.

