## Extended Data and Methods for "Parkinson’s Disease-linked LRRK2 structure and model for microtubule interaction"

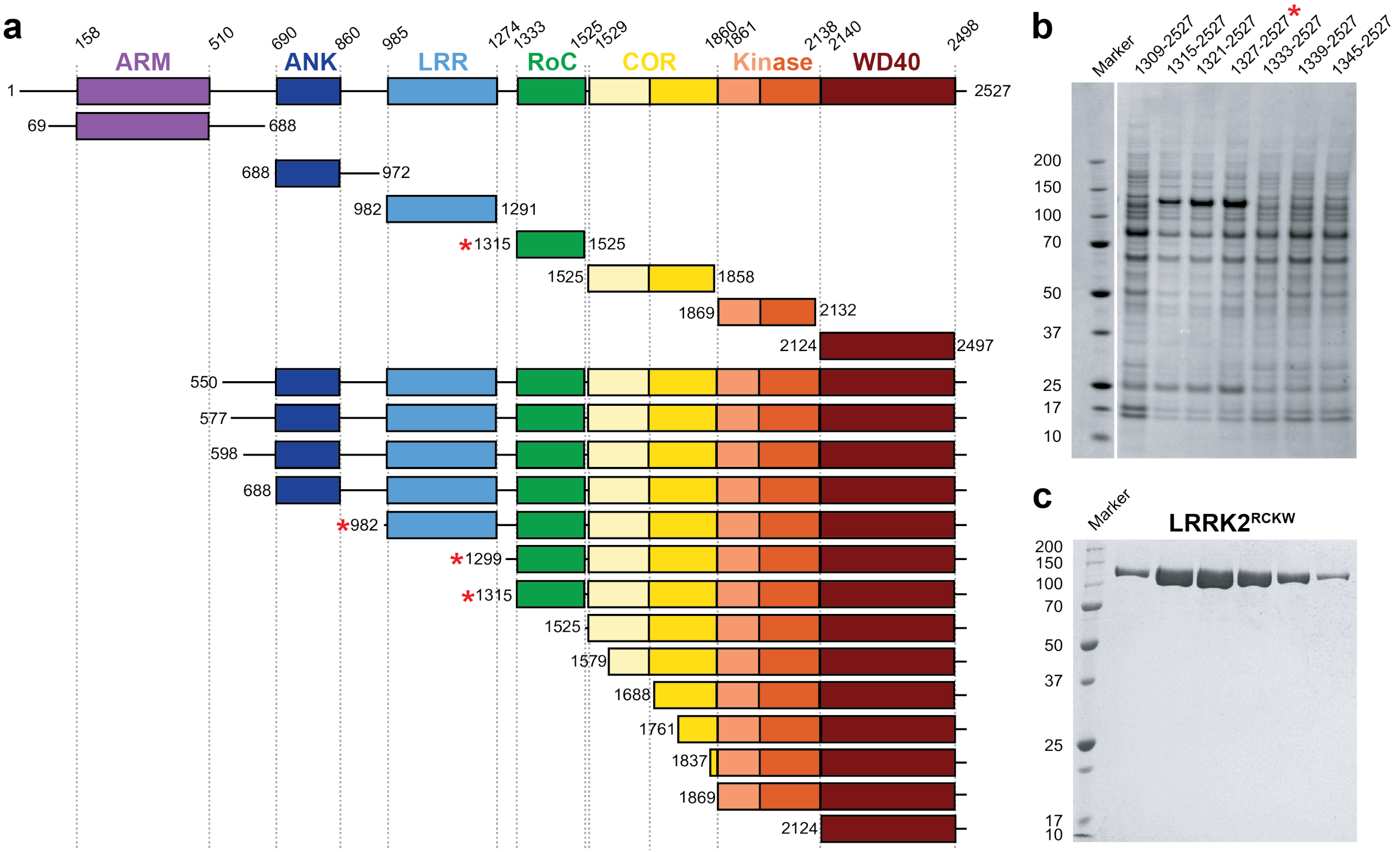


**Extended Data Figure 1 | Optimization of LRRK2 constructs for expression in insect cells. a,** We systematically scanned domain boundaries (amino acid numbers of boundaries noted above domain names) to generate LRRK2 constructs that expressed well in baculovirus-infected insect cells and yielded stable and soluble protein. These attempts included full-length LRRK2, the kinase domain alone or with the WD40 domain, and other isolated domains. In this approach, only the GTPase domain on its own expressed well. Next, we gradually shortened LRRK2 from its amino-terminus. Red asterisks indicate constructs that were soluble. **b,** After identifying domain boundaries yielding constructs that expressed soluble protein, additional fine tuning of boundaries was performed. A Coomassie stained SDS-PAGE gel shows systematic N-terminal truncations at the RoC domain resulting in the identification of a construct with the highest expression levels: amino acids 1327 to 2527 (red asterisk, "LRRK2$^{RCKW}$" here). **c,** A Coomassie stained SDS-PAGE gel of purified LRRK2$^{RCKW}$ after elution from an S200 gel filtration column. As predicted by its primary structure, LRRK2$^{RCKW}$ runs at ~140 kDa.

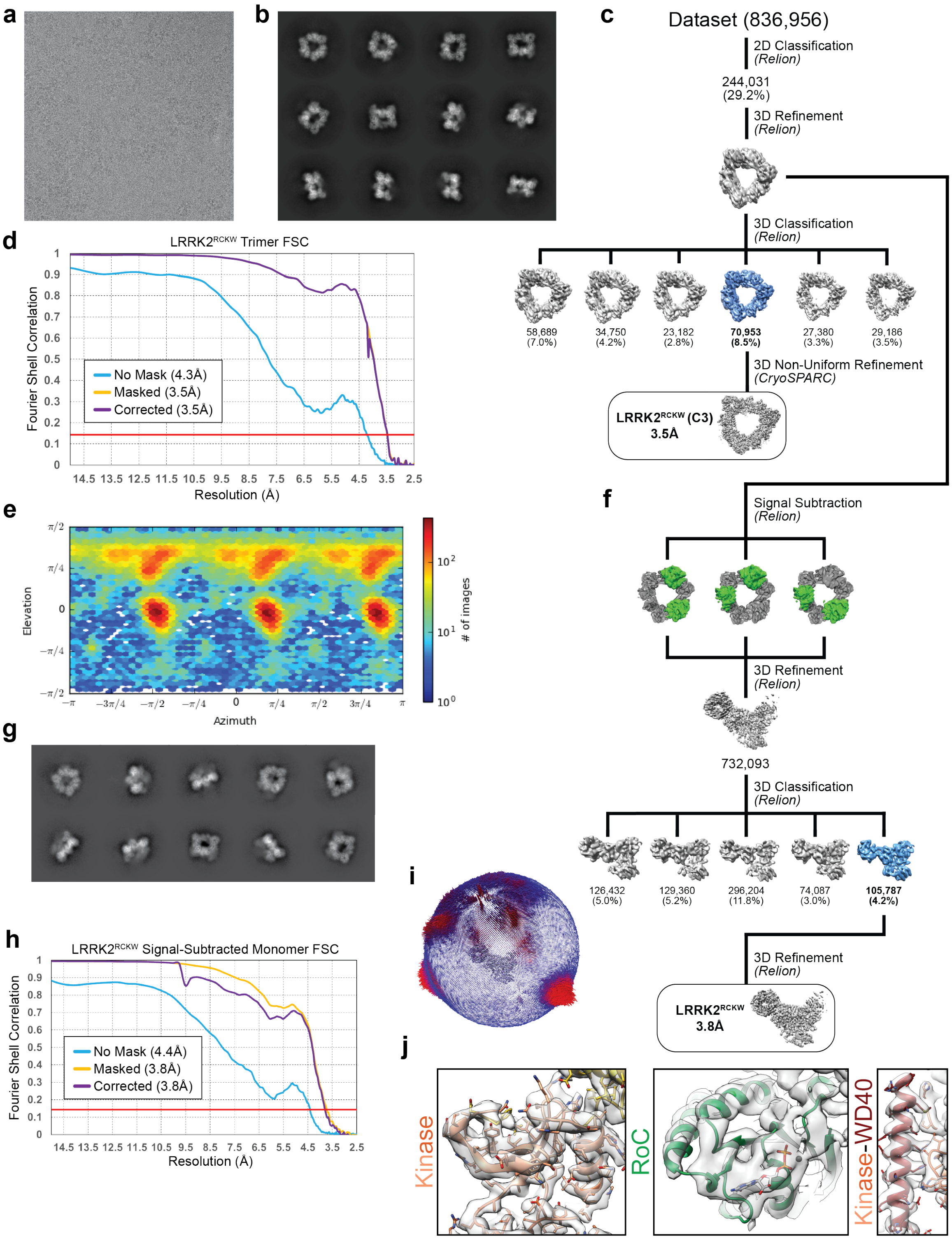


**Extended Data Figure 2 | Cryo-EM structure determination of LRRK2$^{RCKW}$.** **a**, Electron micrograph of LRRK2$^{RCKW}$. **b**, 2D class averages of the LRRK2$^{RCKW}$ trimer. **c**, 2D/3D classification scheme used to obtain the 3.5Å structure of the LRRK2$^{RCKW}$ trimer. **d, e,** Fourier Shell Correlations (from Cryosparc) (d) and Euler angle distribution (e) for the LRRK2$^{RCKW}$ trimer. **f**, Processing strategy used to obtain a 3.8Å structure of LRRK2$^{RCKW}$ generated from a signal-subtracted trimer where only one monomer contains the RoC and COR-A domains. This structure improved the resolution of the RoC and COR-A domains relative to the trimer shown in (c). **g-i**, 2D class averages (g), Fourier Shell Correlations (from Relion) (h), and Euler angle distribution (from Relion) (i) for the 3.8Å-resolution signal-subtracted LRRK2$^{RCKW}$ structure. **j,** Close-ups of different parts of the final structure, with the 3.8Å map as a semi-transparent surface and the model shown using the coloring scheme introduced in Fig. 1a.

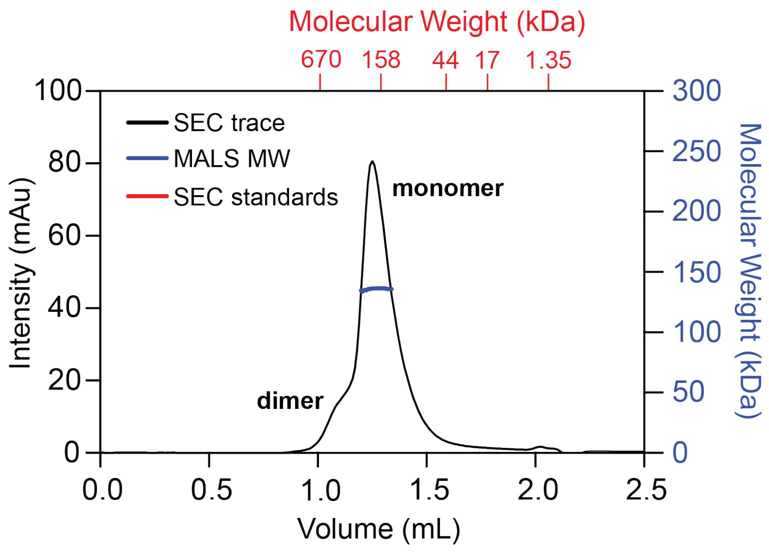


| MW | SEC | MALS |
|---|---|---|
| 137.9kDa | 227kDa | 136kDa |

**Extended Data Figure 3 | LRRK2$^{RCKW}$ is predominantly a monomer under the conditions used for cryo-EM.** Size Exclusion Chromatography-Multiple Angle Light Scattering (SEC-MALS) analysis of LRRK2$^{RCKW}$ under the conditions used for cryo-EM (Fig. 1). The table below the elution profile shows the calculated molecular weights (MW) of LRRK2$^{RCKW}$ according to SEC standards ("SEC") and MALS.

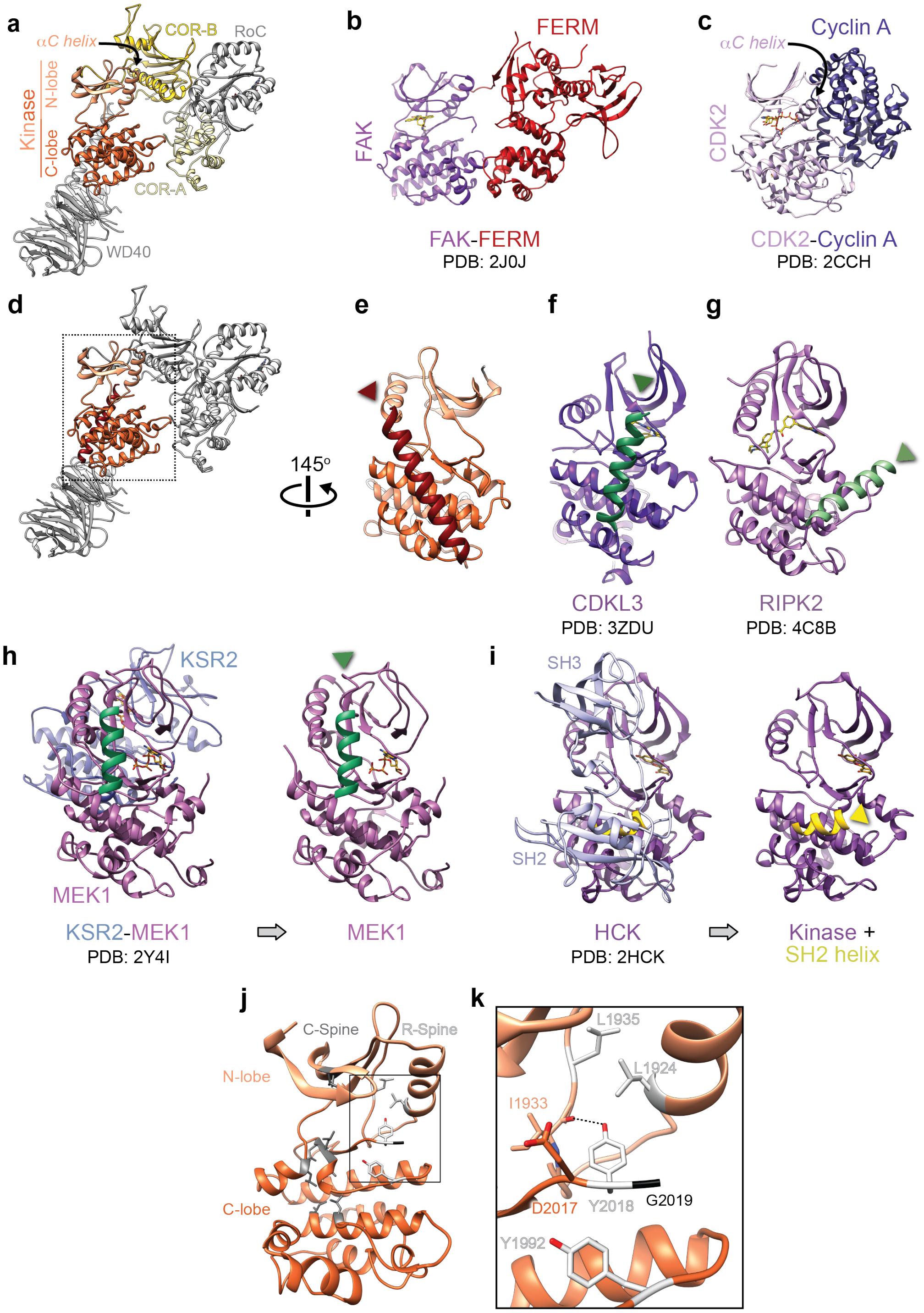


**Extended Data Figure 4 | Comparisons between LRRK2 and other kinases. a**, View of the LRRK2$^{RCKW}$ atomic model with COR-A, COR-B and kinase domains colored. The N- and C-lobes of the kinase are labeled, as is the αC helix in the N-lobe. **b, c,** The FAK-FERM (PDB: 2J0J)[27] (b) and CDK2-Cyclin A (PDB: 2CCH)[29] (c) complexes, shown in the same orientation as the kinase in (a). The αC helix of CDK2 is also labeled. **d,** Same view as in (a) with only the kinase domain and the C-terminal helix colored. **e,** Rotated view of LRRK2's kinase domain with the C-terminal helix facing the viewer. **f, g,** CDKL3 (PDB: 3ZDU) (f) and RIPK2 (PDB: 4C8B)[35] (g) shown in the same orientation as LRRK2's kinase in (e), with alpha helices with the same general location as LRRK2's C-terminal helix colored in green. **h,** KSR2-MEK1 complex (PDB: 2Y4I), with the kinase oriented as in (e) (left) and after removing KSR2 for clarity (right). The alpha helix associated with the kinase is shown in green. **i,** HCK (PDB: 2HCK) in complex with its SH2 and SH3 domains with the kinase oriented as in (e) (left), and after removal of the SH2 and SH3 domains for clarity (right). A remaining alpha helix from the SH2 domain is shown in yellow. **j,** Front view of LRRK2's kinase with the C-Spine and R-Spine residues shown and colored in grey and white, respectively. **k,** Close-up of the DYG motif and neighboring R-Spine residues. I1933, whose backbone carbonyl is within hydrogen-bonding distance (2.7Å) of Y2018, is shown in this panel as well.

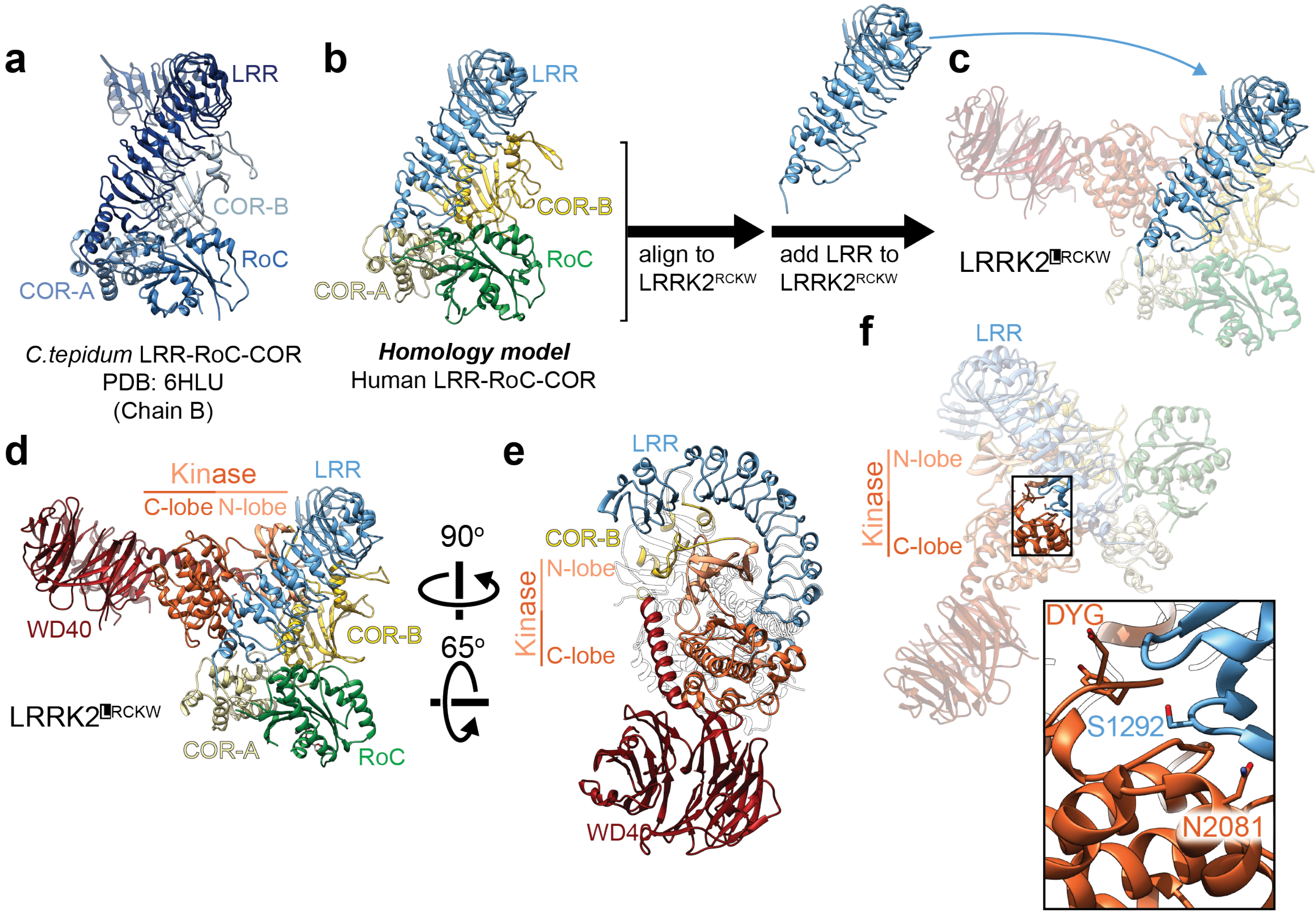


**Extended Data Figure 5 | Modelling of the Leucine-Rich Repeat (LRR) into LRRK2$^{RCKW}$.** **a**, Crystal structure of the LRR-RoC-COR(A/B) domains from *C. tepidum* Roco (PDB: 6HLU)[15]. **b**, Homology model for human LRR-RoC-COR(A/B) based on the *C. tepidum* Roco structure (from SWISS-MODEL). **c**, Chimeric model combining LRRK2$^{RCKW}$ and the homology model for the LRR domain from (b) obtained by aligning their RoC-COR(A/B) domains. **d**, **e**, Two views of the hybrid LRRK2$^{LRCKW}$ model. **f**, Close-up showing the proximity between the active site of the kinase (with the side chains of its DYG motif shown) and the S1292 autophosphorylation site on the LRR. The close-up also highlights the proximity between N2081, a residue implicated in Crohn's Disease, and the LRR.

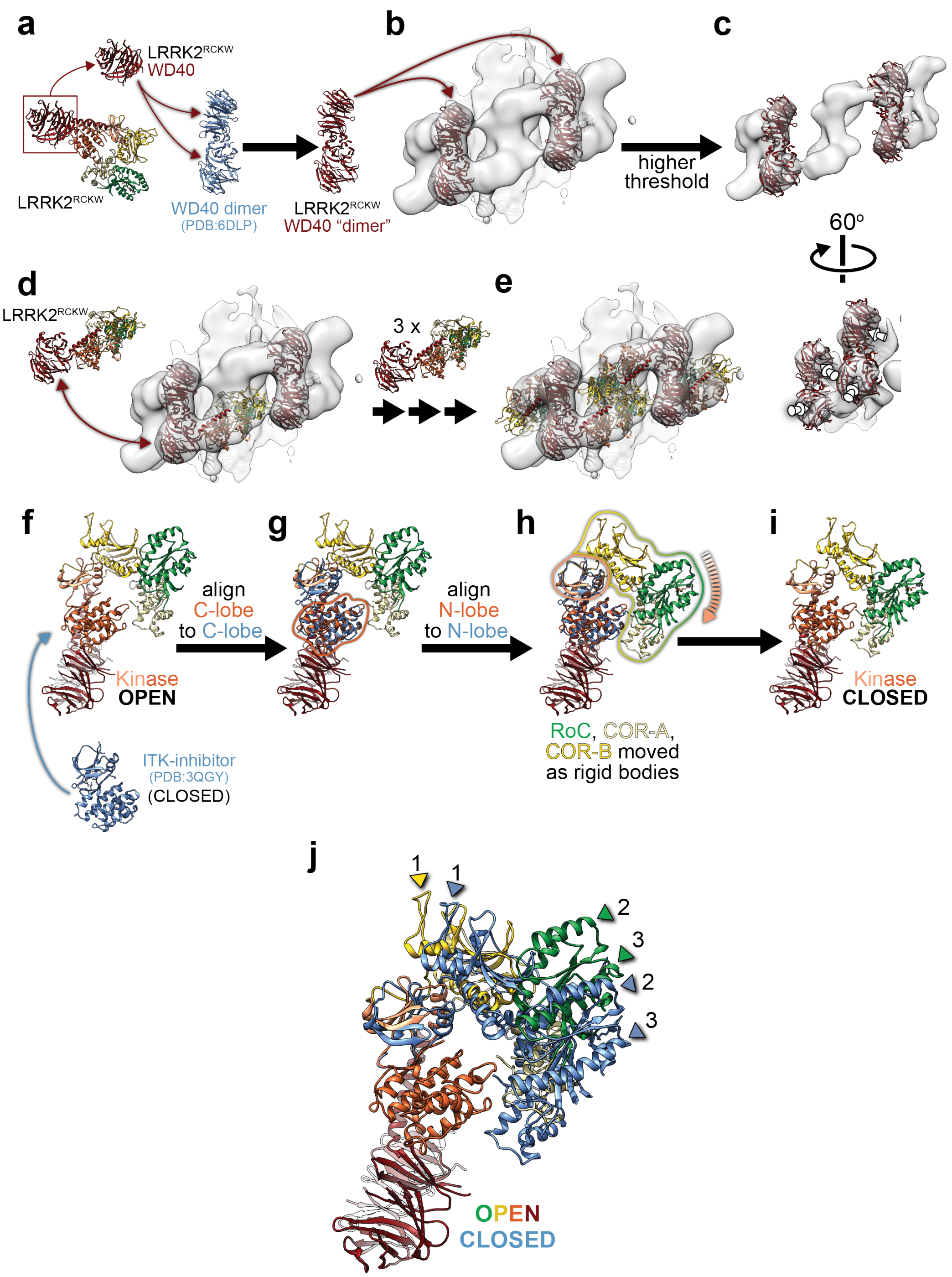


**Extended Data Figure 6 | Docking of LRRK2$^{RCKW}$ into the sub-tomogram average of cellular LRRK2 filaments, and modeling of the closed-kinase LRRK2$^{RCKW}$ filaments.** **a**, The WD40s in the crystal structure of a dimer of LRRK2's WD40 (PDB: 6DLP)[18] were replaced with the WD40s from our cryo-EM structure of LRRK2$^{RCKW}$. **b,** The resulting dimer was fitted into the 14Å sub-tomogram average of cellular microtubule-associated LRRK2 filaments. **c,** Two views of the same fitting shown in (b), displayed with a higher threshold for the map to highlight the fitting of the WD40 ß-propellers into the density. **d**, Four copies of LRRK2$^{RCKW}$ were docked into the sub-tomogram average by aligning their WD40 domains to the docked WD40 dimer. **e**, Model containing the four aligned LRRK2$^{RCKW}$. **f-i**, Modeling of the kinase-closed form of LRRK2$^{RCKW}$. **f, g,** The structure of ITK bound to an inhibitor (PDB: 3QGY)[54], which is in a closed conformation, was aligned to LRRK2$^{RCKW}$ using only the C-lobes of the two kinases. **h**, The N-terminal portion of LRRK2$^{RCKW}$, comprising RoC, COR-A, COR-B and the N-lobe of the kinase, was aligned to ITK using only the N-lobes of the kinases. RoC, COR-A and COR-B were moved as a rigid body in this alignment. **i**, A kinase-closed model of LRRK2$^{RCKW}$. **j**, Superposition of the kinase-open (our cryo-EM structure) and kinase-closed LRRK2$^{RCKW}$ (the model built here, in blue) showing the closing of the RoC-COR(A/B)-kinase N-lobe portion of LRRK2$^{RCKW}$. Three structural reference points are shown in colors and numbered to highlight the differences between the two structures.

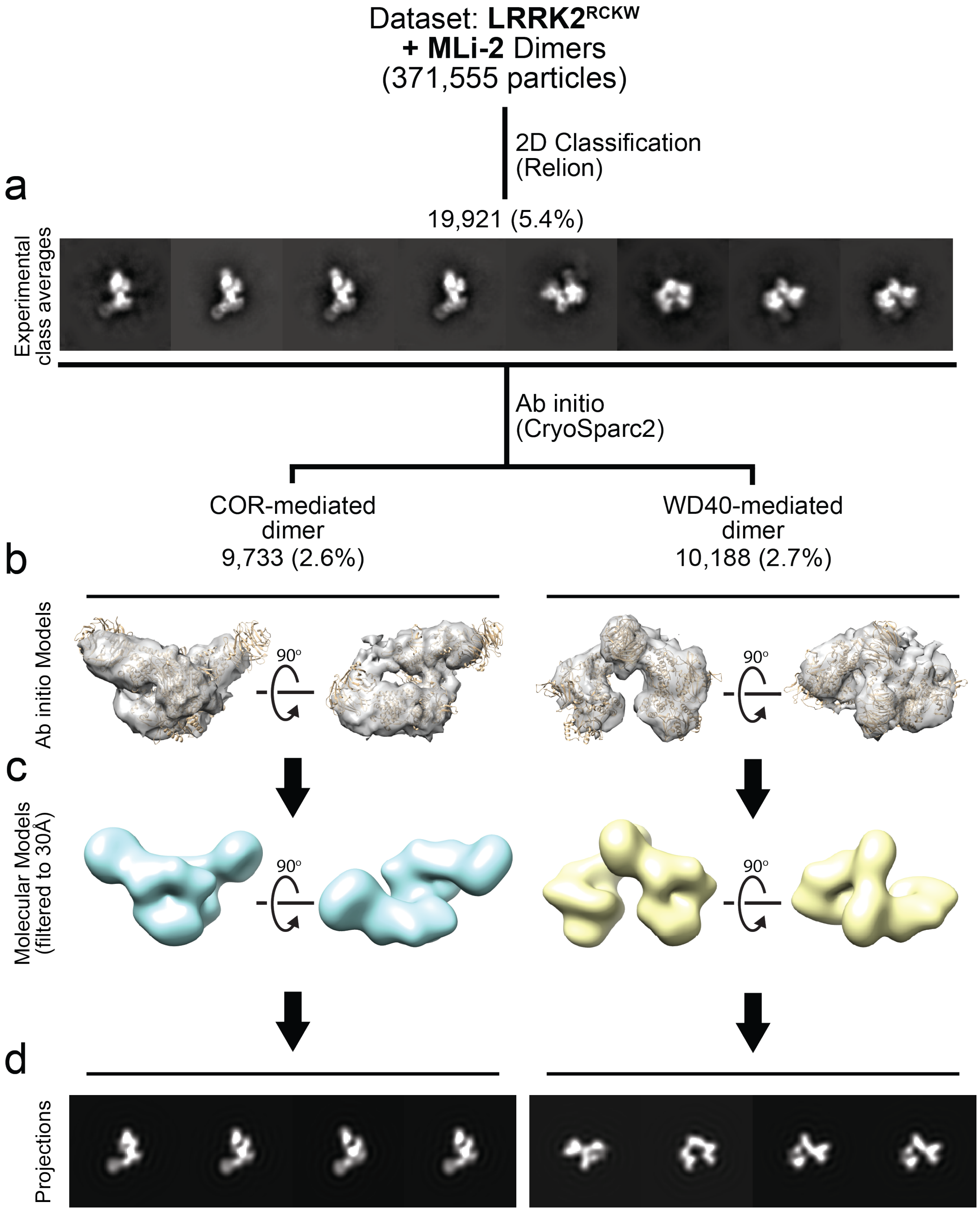


**Extended Data Figure 7 | Generation of *ab initio* models for cryo-EM of LRRK2$^{RCKW}$ dimers.** An initial dataset was collected from a sample of LRRK2$^{RCKW}$ incubated in the presence of the kinase inhibitor MLi-2 and dimers were selected. **a,** Representative two-dimensional class averages used for *ab initio* model building. **b,** *Ab initio* models with the structure of LRRK2$^{RCKW}$ docked in. **c,** Volumes generated form the molecular models in (b), filtered to 30Å resolution. **d,** Projections of the volumes in (c) shown in the same order as their corresponding 2D class averages in (a).

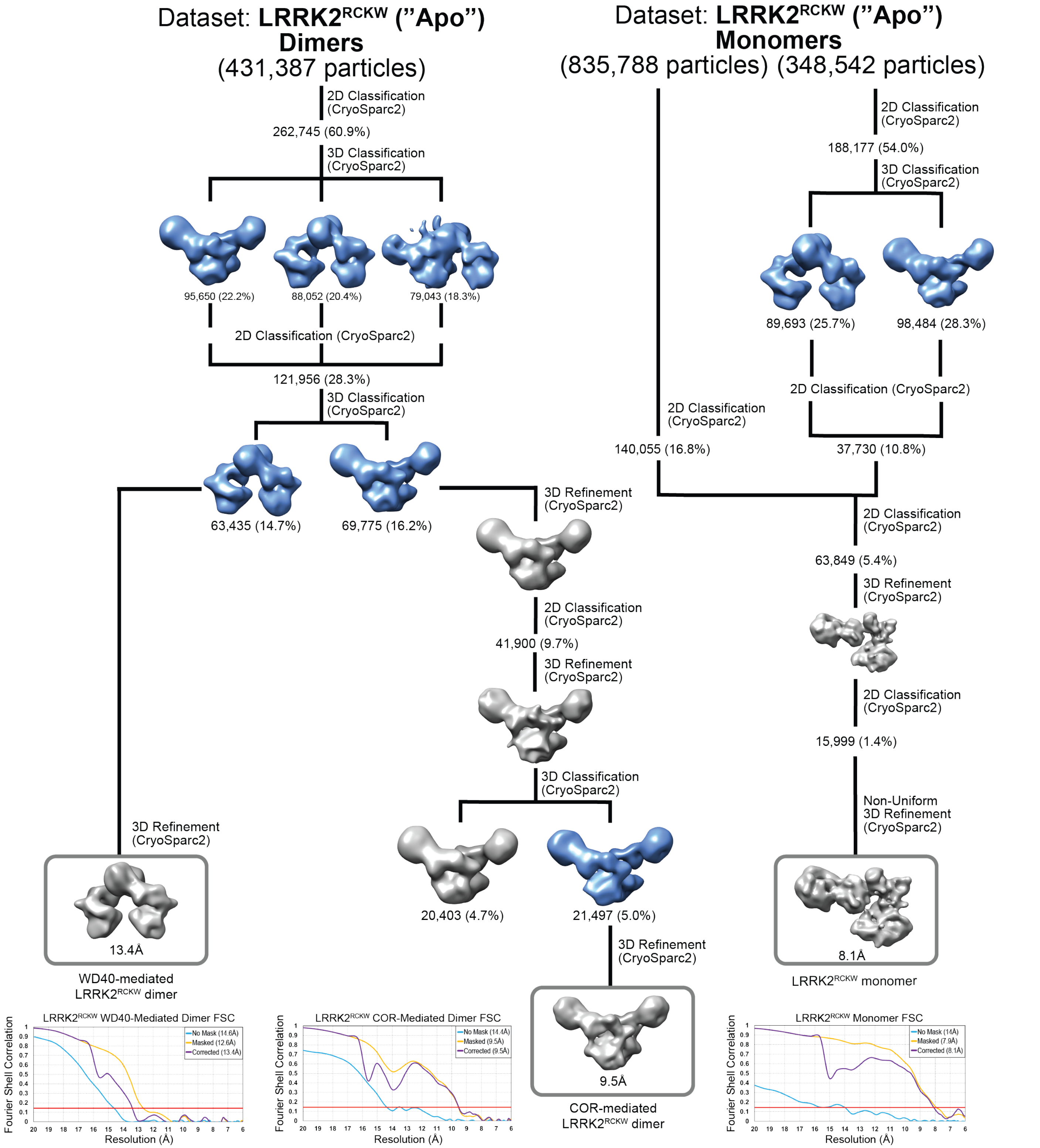


**Extended Data Figure 8 | Data processing strategy for obtaining cryo-EM structures of a monomer and WD40- and COR-mediated dimers of LRRK2$^{RCKW}$ in the absence of inhibitor ("Apo").** The models used during the processing of the dimers (see Methods) are those shown in Extended Data Fig. 7 along with an additional linear trimer (see Methods) used for particle sorting. The models used for processing of the monomer (see Methods) were the same dimer models as in Extended Data Fig. 7 (used for particle sorting) in addition to a monomer model generated from our LRRK2$^{RCKW}$ model (used for refinement).

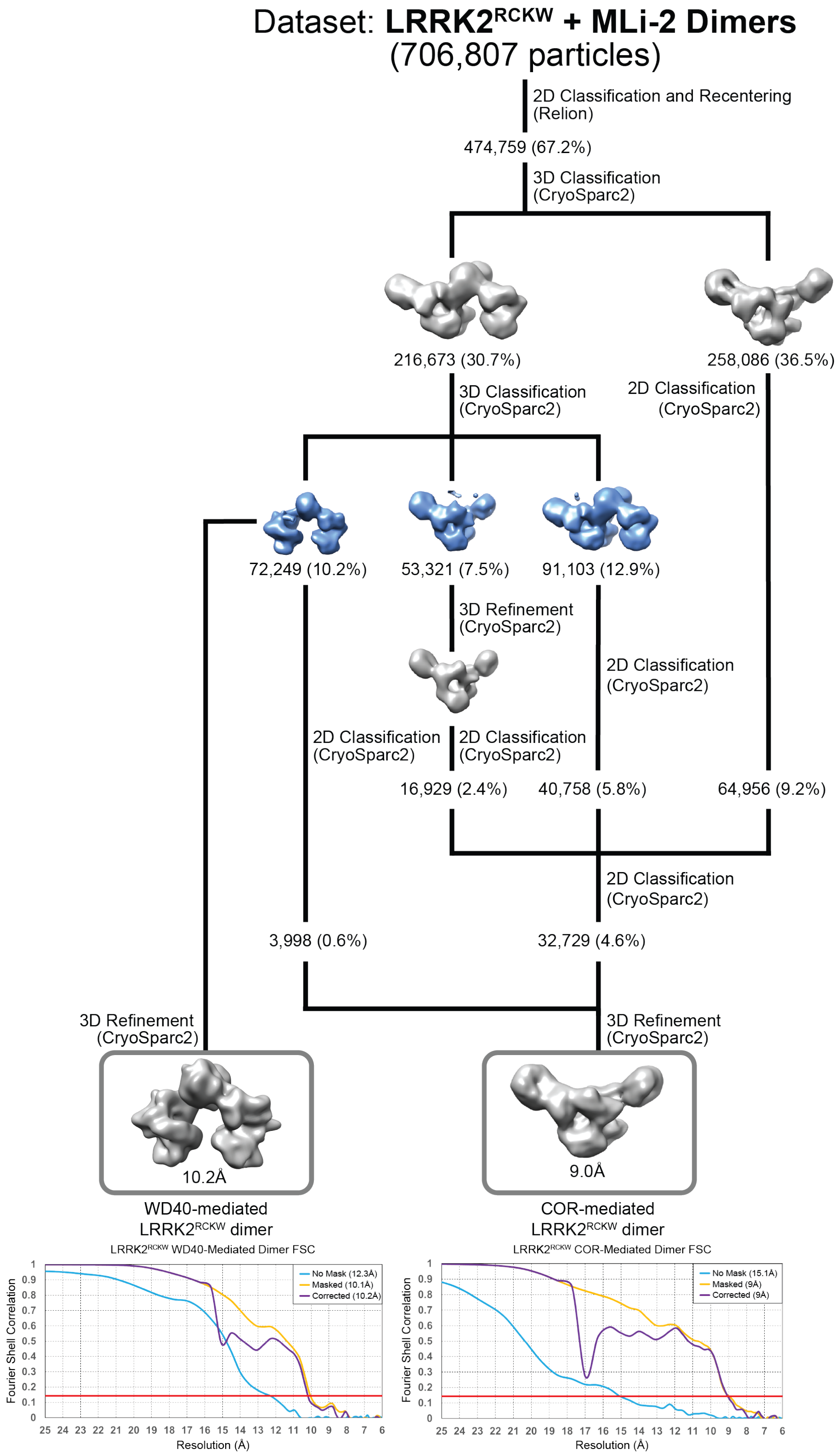


**Extended Data Figure 9 | Data processing strategy for obtaining cryo-EM structures of WD40- and COR-mediated dimers of LRRK2$^{RCKW}$ in the presence of the inhibitor MLi-2.** The models used during this processing (see Methods) are those shown in Extended Data Fig.7 along with an additional linear trimer (see Methods) used for particle sorting.

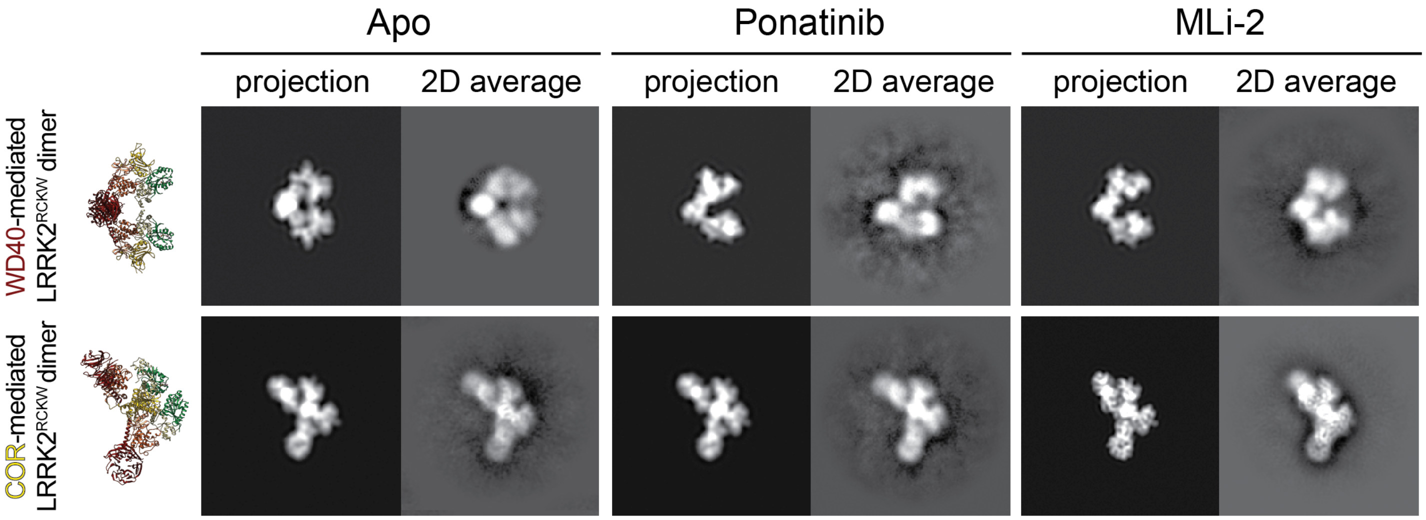


**Extended Data Figure 10 | LRRK2$^{RCKW}$ forms WD40- and COR-mediated dimers outside the filaments.** Two-dimensional (2D) class averages of WD40- and COR-mediated LRRK2$^{RCKW}$ dimers obtained in the absence of inhibitors ("Apo") or in the presence of either Ponatinib or MLi-2. The same molecular models of the two dimers shown in Fig. 3 are shown on the left but in orientations similar to those represented by the 2D class averages shown here. For each class average, a projection from the corresponding model in the best-matching orientation is shown to its left.

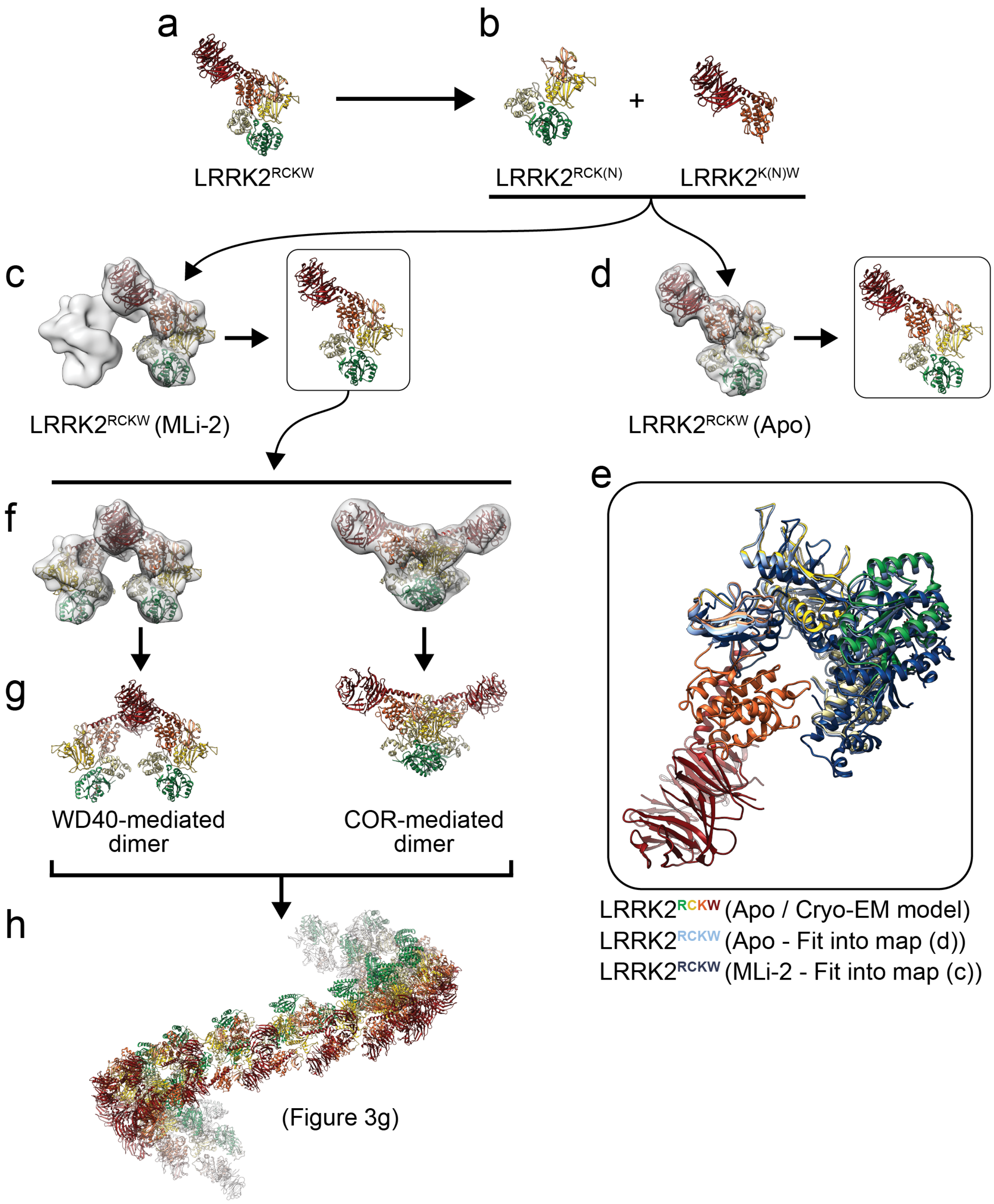


**Extended Data Figure 11 | The basic geometric properties of the microtubule-associated LRRK2$^{RCKW}$ filaments are encoded in the structure of LRRK2$^{RCKW}$. a, b,** The LRRK2$^{RCKW}$ structure solved in this work (a) was split at the junction between the N- and C-lobes of the kinase domain (L1949-A1950) (b). **c, d,** Docking of the two halves of LRRK2$^{RCKW}$ into cryo-EM maps of LRRK2$^{RCKW}$ solved in the presence of MLi-2 (c) or without inhibitor ("Apo") (d). The dimer maps are the same ones shown in Fig. 3 and Extended Data Figs. 8 and 9 and the Apo map is the one shown in Fig. 1g, h and Extended Data Fig. 8. **e**, Three-way comparison of LRRK2$^{RCKW}$ (with domain colors) and the models resulting from the dockings into the MLi-2 WD40-mediated dimer map (c) (dark blue) and "Apo" monomer map (d) (light blue). The three structures were aligned using the C-lobes of their kinases and the WD40 domain. The superposition illustrates that the docking into the "Apo" map results in a structure very similar to that obtained from the trimer (Fig. 1) and that the presence of MLi-2 leads to a closing of the kinase. **f,** The model obtained in (c) was docked into cryo-EM maps of either WD40- or COR-mediated dimers obtained in the presence of MLi-2. **g,** Molecular models resulting from the docking in (f). **h**, Aligning, in alternating order, copies of the dimer models generated in (f, g) results in a right-handed filament with dimensions compatible with those of a microtubule, and its RoC domains pointing inwards (see Fig. 3g-i for more details).

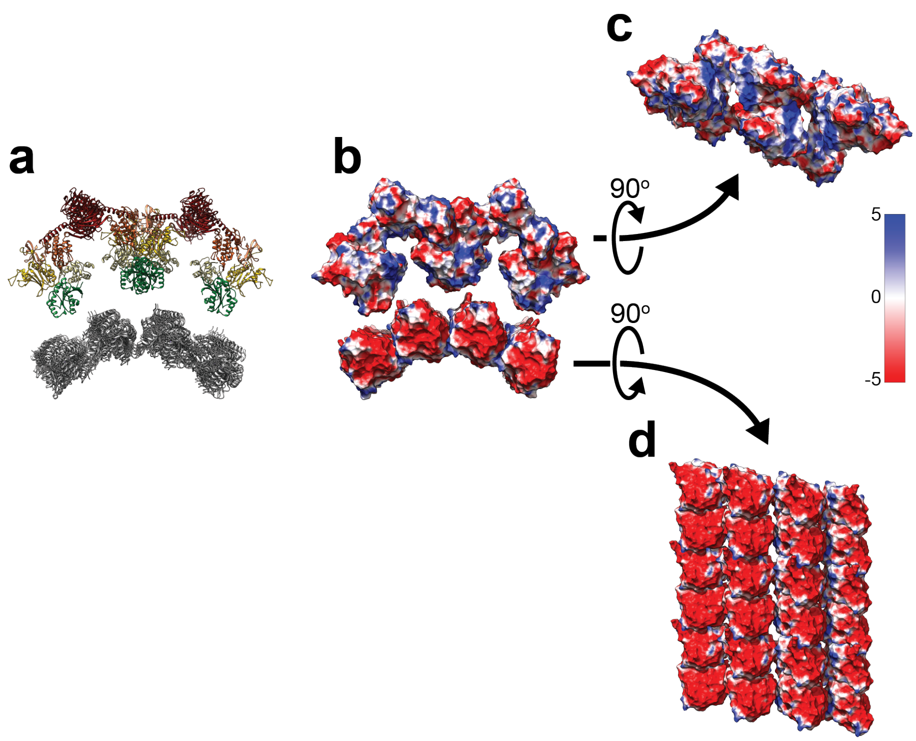


**Extended Data Figure 12 | Complementarity between the surface charge distributions of the LRRK2$^{RCKW}$ filament model and the microtubule. a**, Molecular model of the microtubule-associated LRRK2$^{RCKW}$ filament obtained by docking a fragment of a microtubule structure (PDB: 6O2S) into the corresponding density in the sub-tomogram average (Fig. 2a). **b**, Same view as in (a) with the models shown as surface representations colored by their Coulomb potential. **c, d,** "Peeling off" of the structure shown in (b), with the LRRK2$^{RCKW}$ filament seen from the perspective of the microtubule surface (c) and the microtubule surface seen from the perspective of the LRRK2$^{RCKW}$ filament (d). Note: the acidic C-terminal tubulin tails are not ordered in the microtubule structure and thus are not included in the surface charge distributions. The Coulomb potential coloring scale is shown on the right.

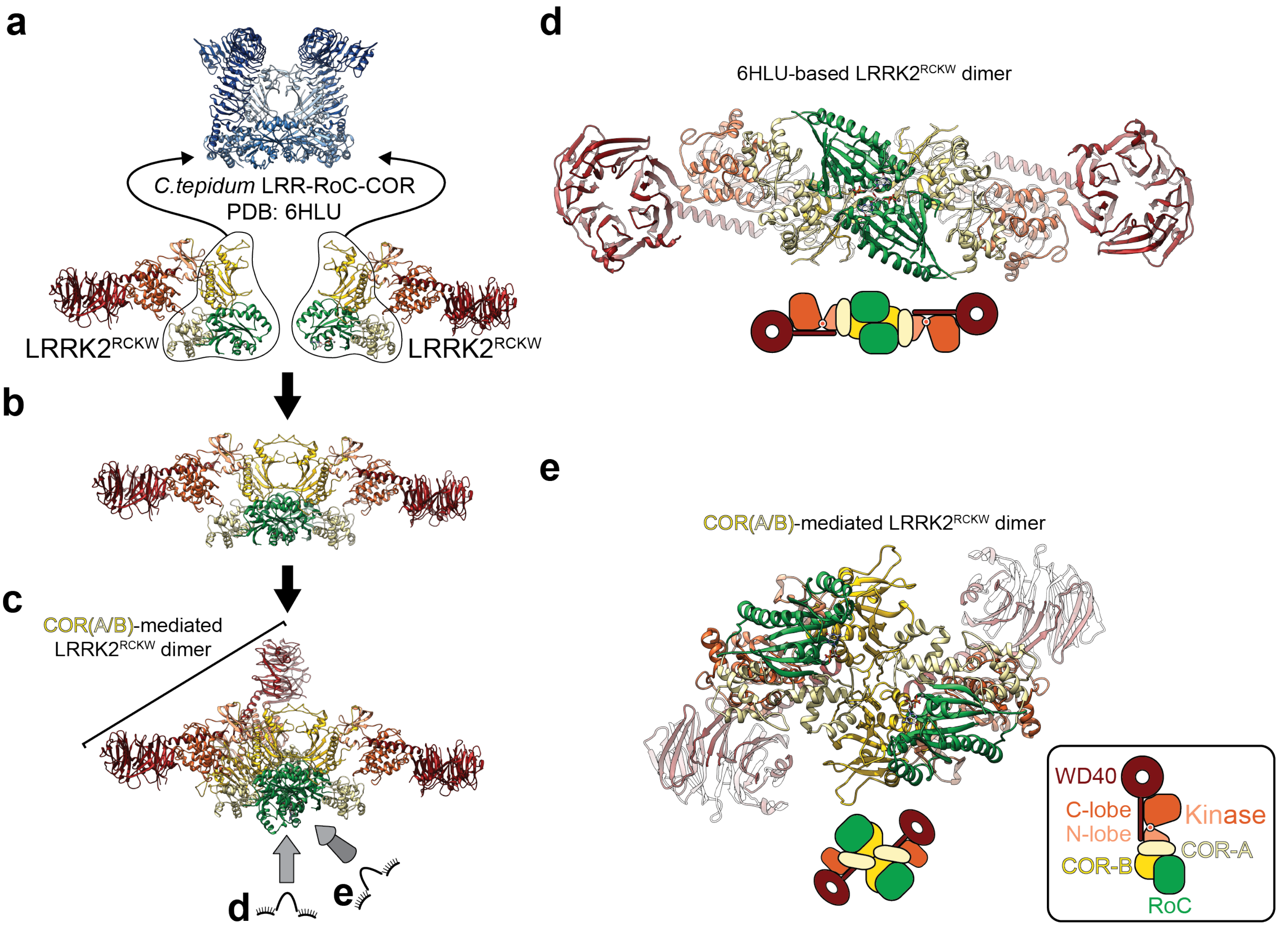


**Extended Data Figure 13 | LRRK2$^{RCKW}$'s COR-mediated dimerization interface differs from that of the *C. tepidum*'s Roco homolog. a, b**, Two copies of the LRRK2$^{RCKW}$ structure were aligned to the RoC-COR domains of the LRR-RoC-COR structure from *C. tepidum*'s Roco protein (PDB: 6HLU) (a) to replicate the interface observed in the bacterial homolog in the context of the human protein (b). **c**, One of the LRRK2$^{RCKW}$ monomers was used to superimpose the dimer modeled in (b) with the COR-mediated dimer seen both in the microtubule-associated filaments in cells (cryo-ET) (Fig. 2) as well as in the absence of microtubules (by cryo-EM) (Fig. 3). **d**, **e**, Comparison between the dimer modeled based on the *C. tepidum* LRR-RoC-COR structure (d) and the dimer observed for LRRK2$^{RCKW}$ in this work (e). While the bacterial structure shows a dimerization interface that involves the GTPase (RoC), LRRK2$^{RCKW}$ interacts exclusively through its COR-A and -B domains, with the RoC domains located away from this interface. The two arrangements are shown schematically in cartoon form below the structures.

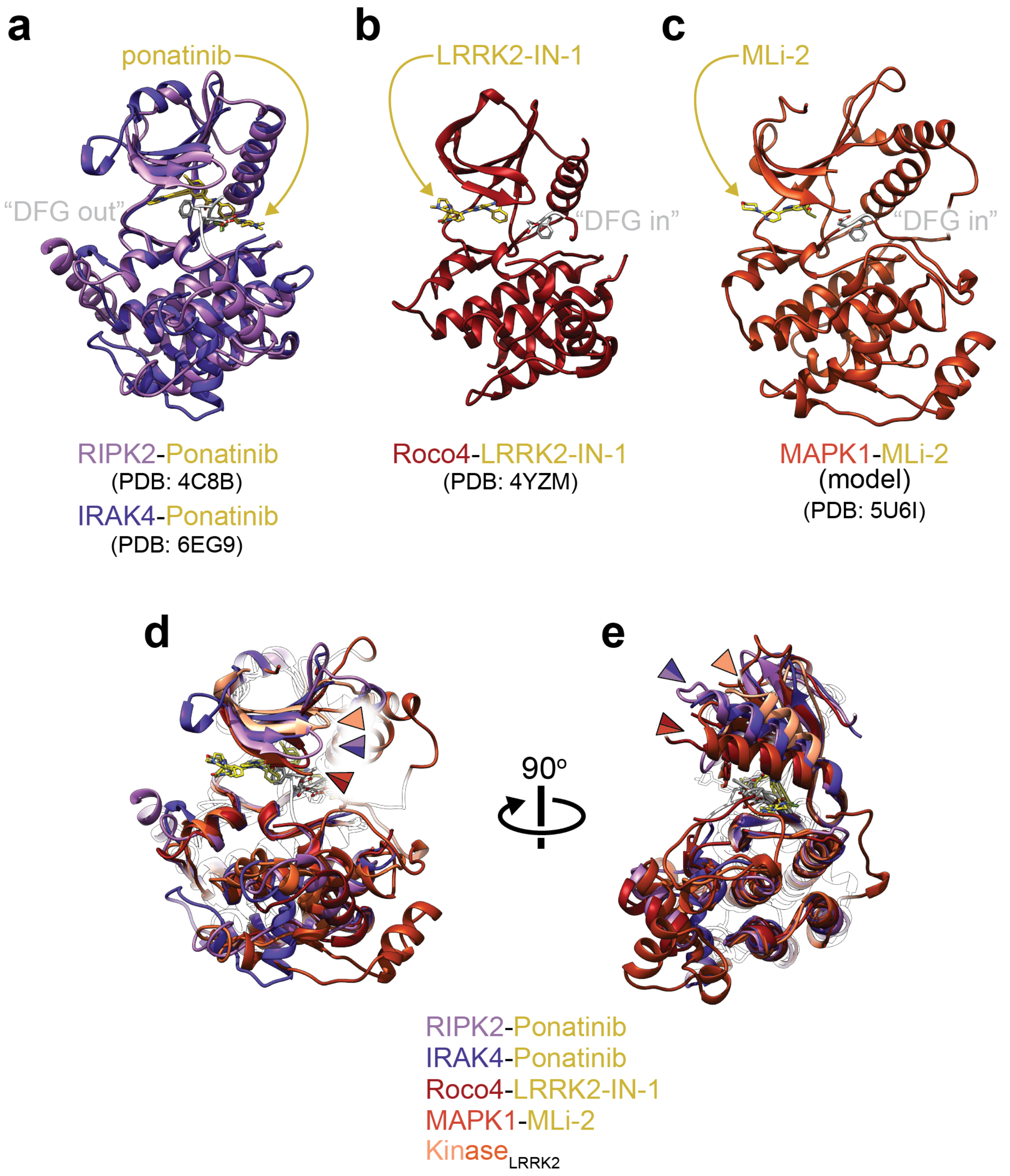


**Extended Data Figure 14 | Ponatinib is a Type 2, "DFG out" inhibitor. a**, Superposition of the structures of Ponatinib-bound RIPK2 (PDB: 4C8B)[55] and IRAK4 (PDB: 6EG9). Ponatinib is shown in yellow, and the DYG motif residues are shown in white. **b**, **c** For comparison, the structures of (a) Roco4 bound to LRRK2-IN-1 (PDB: 4YZM), a LRRK2-specific Type 1, "DFG in" inhibitor, and (b) a model of Mitogen-activated kinase 1 (MAPK1) bound to MLi-2 (PDB: 5U6I), another LRRK2-specific Type 1, "DFG in" inhibitor are shown. The inhibitor and DFG residues are colored as in (a). **d**, The structures in (a-c), as well as the kinase from LRRK2$^{RCKW}$ are shown superimposed. The color arrowheads point to the N-lobe's ß-sheet to highlight the difference in conformation between kinases bound to the two different types of inhibitors. Note that LRRK2$^{RCKW}$'s kinase is even more open than the two Ponatinib-bound kinases. **e**, Rotated view of (d), now highlighting the position of the N-lobe's $\alpha$C helix. An additional alpha helix in the N-lobe of MAPK1 was removed from this view for clarity.

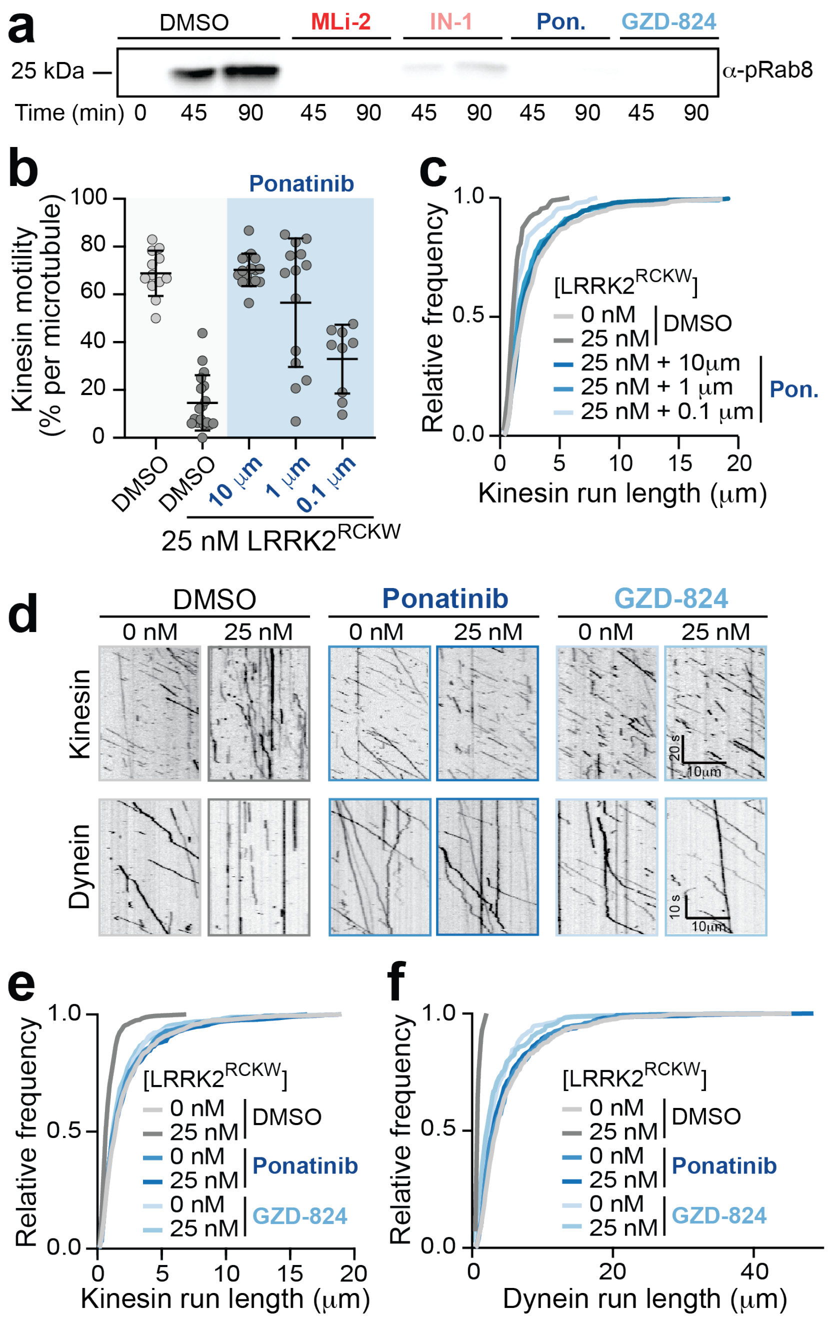


**Extended Data Figure 15 | Type 2 kinase inhibitors rescue microtubule-based motor motility. a,** The kinase inhibitors MLi-2 (1 μM), LRRK2-IN-1 (1 μM), Ponatinib (10 μM) and GZD-824 (10 μM) all inhibit LRRK2$^{RCKW}$'s kinase activity *in vitro* compared to a DMSO control. A Western blot using a phospho-specific antibody to Rab8a at the indicated time points is shown. **b,** A dose response curve showing the percentage of motile kinesin events per microtubule as a function of Ponatinib concentration with LRRK2$^{RCKW}$ (25 nM) or without LRRK2$^{RCKW}$. Data are mean ± s.d. (from left to right: n = 12, 18, 16, 14, and 9 microtubules quantified from one experiment). ****p < 0.0001 calculated using the Kruskal-Wallis test with Dunn's posthoc for multiple comparisons (compared to DMSO without LRRK2$^{RCKW}$). **c,** Dose response curve of run lengths from data in (b) represented as a cumulative frequency distribution. From top to bottom: n = 654, 173, 584, 293, and 129 motile kinesin events. Mean decay constants (tau) ± confidence interval (CI) are (from top to bottom) 2.736 ± 0.113, 1.291 ± 0.181, 2.542 ± 0.124, 2.285 ± 0.134, and 1.653 ± 0.17. **d,** Representative kymographs of kinesin and dynein with DMSO or Type 2 inhibitors with or without LRRK2$^{RCKW}$. **e,** The Type 2 kinase inhibitors Ponatinib and GZD-824 rescue kinesin run length, represented as a cumulative frequency distribution of run lengths with LRRK2$^{RCKW}$ (25 nM) or without LRRK2$^{RCKW}$ (0 nM). From top to bottom: n = 893, 355, 507, 499, 524, and 529 runs from two independent experiments. Mean decay constants (tau) ± 95% CI are (from top to bottom) 2.070 ± 0.058, 0.8466 ± 0.091, 1.938 ± 0.065, 2.075 ± 0.07, 1.898 ±, 0.065, and 1.718 ± 0.064. **f,** Same as in (e) but with dynein. From top to bottom: n = 659, 28, 289, 306, 254, and 339 runs from two independent experiments). Mean decay constants (tau) ± 95% confidence intervals; microns are 4.980 ± 0.147, 0.846 ± 0.415, 4.686 ± 0.142, 4.445 ± 0.172, 3.156 ± 0.09, 3.432 ± 0.188 (from top to bottom). The DMSO conditions are reproduced from Fig. 4f for comparison. See Extended Data Table 1 for all source data and replicate information.

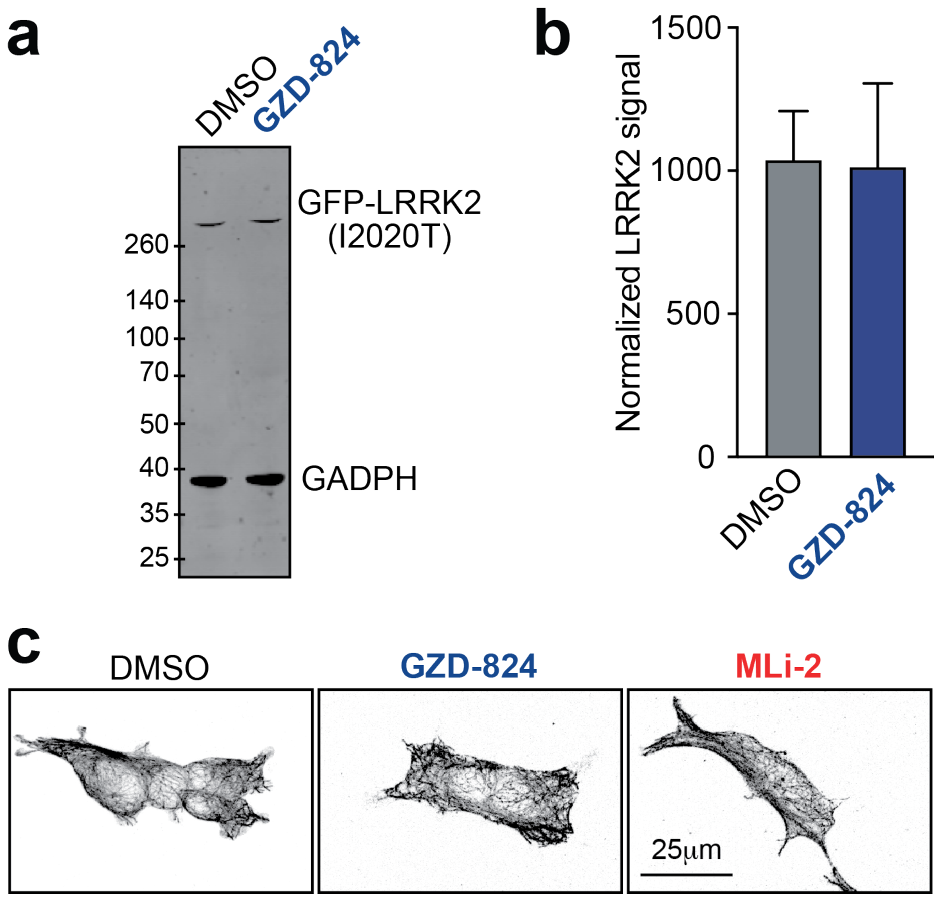


**Figure 16 | Microtubule architecture and LRRK2 expression are not perturbed by kinase inhibitors**. **a,** Expression levels of GFP-LRRK2 (I2020T) in 293T cells treated with either DMSO or GZD-824 (5 µM). An Immunoblot with anti-GFP (LRRK2) and anti-GADPH (loading control), which is a representative image from three replicates, is shown. **b,** Quantification of GFP-LRRK2 (I2020T) expression levels from Western blots similar to (a). Data are mean ± s.d. (n = 3 per condition). GZD-824 is not significantly different from the DMSO-treated control (Mann-Whitney test). **c,** 293T cells immunostained for tubulin showing that the microtubule architecture is not affected by GZD-824 or MLi-2 compared to DMSO treatment. See Extended Data Table 1 for all source data and replicate information.

### Materials and Methods

Cloning, plasmid construction, and mutagenesis

For baculovirus expression, the DNA coding for the LRRK2 residues 1327 to 2527 (taken from Mammalian Gene Collection) was PCR-amplified using the forward primer TACTTCCAATCCATGAAAAAGGCTGTGCCTTATAACCGA and the reverse primer TATCCACCTTTACTGTCACTCAACAGATGTTCGTCTCATTTTTTCA. The T4 polymerase-treated amplicon was inserted into the expression vector pFB-6HZB by ligation-independent cloning. According to Bac-to-Bac expression system protocols (Invitrogen), this plasmid was used for the generation of recombinant Baculoviruses.

For mammalian expression vectors, pDEST53-GFP-LRRK2 (WT)[56] from Addgene (#25044) was used. pDEST53-GFP-LRRK2 (I2020T) was cloned using QuikChange site-directed mutagenesis (Agilent) with the forward primer AAGATTGCTGACTACGGCACTGCTCAGTACTGCTG and the reverse primer CAGCAGTACTGAGCAGTGCCGTAGTCAGCAATCTT. pET17b-Kif5b(1-560)-GFP-His[57] was obtained from Addgene (#15219). For pET28a-ZZ-TEV-Halo-NINL$^{1-702}$, Ninein-like$^{1-702}$ (NINL) was synthesized as previously described[33] and inserted into a pET28a expression vector with a synthesized ZZ-TEV-Halo gBlock fragment (IDT) using Gibson assembly.

LRRK2$^{RCKW}$ expression and purification

The expression construct contained an N-terminal $His_6$-Z-tag, cleavable with TEV protease (Extended Data Fig. 1). For LRRK2$^{RCKW}$ purification, the pelleted Sf9 cells were washed with PBS, resuspended in lysis buffer (50 mM HEPES pH 7.4, 500 mM NaCl, 20 mM imidazole, 0.5 mM TCEP, 5% glycerol, 5 mM $MgCl_2$, 20 µM GDP) and lysed by sonication. The lysate was cleared by centrifugation and loaded onto a Ni-NTA (Qiagen) column. After vigorous rinsing with lysis buffer the $His_6$-Z-tagged protein was eluted in lysis buffer containing 300 mM imidazole. Immediately thereafter, the eluate was diluted with a buffer containing no NaCl, in order to reduce the NaCl-concentration to 250 mM and loaded onto an SP sepharose column. $His_6$-Z-TEV-LRRK2$^{RCKW}$ was eluted with a 250 mM to 2.5 M NaCl gradient and treated with TEV protease overnight to cleave the $His_6$-Z-tag. Contaminating proteins, the cleaved tag, uncleaved protein and TEV protease were removed by another combined SP sepharose Ni-NTA step. Finally, LRRK2$^{RCKW}$ was concentrated and subjected to gel filtration in storage buffer (20 mM HEPES pH 7.4, 800 mM NaCl, 0.5 mM TCEP, 5% glycerol, 2.5 mM $MgCl_2$, 20 µM GDP) using an AKTA Xpress system combined with an S200 gel filtration column. The final yield as calculated from UV absorbance was 1.2 mg LRRK2$^{RCKW}$/L insect cell medium.

SEC-MALS

SEC-MALS experiments were performed using an ÄKTAmicro chromatography system hooked up to a Superdex 200 Increase 3.2/300 size exclusion chromatography column coupled in-line to a DAWN HELEOS II multiangle light scattering detector (Wyatt Technology) and an Optilab T-rEX refractive index detector (Wyatt Technology). SEC-MALS was performed in 50 mM Hepes pH 7.4, 200 mM NaCl, 0.5 mM TCEP, 5% glycerol, 5 mM MgCl2, and 20 µM GDP. For a typical sample, 50 uL of ~7 µM LRRK2$^{RCKW}$ was injected onto the column. Molar mass was calculated using ASTRA 6 software, with protein concentration derived from the Optilab T-rEX. LRRK2$^{RCKW}$ used for SEC-MALS experiments contained an extra 16 residue N-terminal Gly-Ser linker sequence.

Electron microscopy

*Electron microscopy sample preparation and imaging of trimer dataset:*

Purified LRRK2$^{RCKW}$ was dialyzed into a final buffer consisting of 20 mM HEPES pH7.4, 80 mM NaCl, 0.5 mM TCEP, 5% glycerol, 2.5 mM $MgCl_2$ and 20 µM GDP and then diluted to a final concentration of 4 µM in the same buffer. This sample was applied to glow-discharged (20mA for 20s in a K100 Instrument) UltrAuFoil Holey Gold R 1.2/1.3 grids (Quantifoil). A Vitrobot (FEI) was then used to blot away excess sample and plunge freeze the grids in liquid ethane. Grids were stored in liquid nitrogen until imaged.

Cryo-EM data was collected at UCLA California NanoSystems Institute in a Titan Krios (FEI) operated at 300kV, equipped with a K2 Summit direct electron detector (Gatan) and a Quantum energy filter (Gatan). Automated data collection was performed using Leginon[58]. We recorded a total of 3,824 movies in ‘counting mode’ at a dose rate of 6.65 electrons $Å^{-2}$ $sec^{-1}$ with a total exposure time of 8s sub-divided into 200ms frames, for a total of 40 frames. The images were recorded at a nominal magnification of 130,000x resulting in an object pixel size of 1.07Å. The defocus range of the data was -1 µm to -1.8 µm.

*Electron microscopy map and model generation of trimer dataset:*

We aligned the movie frames using UCSF MotionCor2[59], using the dose-weighted frame alignment option. We estimated the CTF on dose-weighted images using GCTF version 1.06[60] as implemented in Appion[61] with per-particle CTF generation. Images having CTF fits worse that 5Å (as determined by GCTF) were excluded from further processing. Using this approach, 3,693 micrographs were kept for further processing. We selected particles from micrographs using FindEM[62] with projections of a trimeric LRRK2$^{RCKW}$ map, created from an initial Cryosparc *ab initio* model generation, serving as a reference. Particle picking was performed within the framework of Appion, resulting in a data set of 836,956 particles.

We carried out subsequent processing first in Relion 3.0[63] then in CryoSparc2[64]. A series of 2D and 3D classifications were performed as shown in Extended Data Fig. 2c to generate the final map. The initial reference was created in a similar manner to those used for template picking, from an initial *ab initio* model generated in Cryosparc. All references were filtered to either 60Å (default in Relion) or 30Å (default in CryoSparc) before refinement processes. The final map, generated in Cryosparc2 using non-uniform refinement and while applying C3 symmetry, reached 3.47Å resolution. Initial 2D classifications used binned data (4.28Å $pixel^{-1}$) while all subsequent 3D classifications and refinement steps used unbinned images (1.07Å $pixel^{-1}$).

In order to improve the density corresponding to the RoC and COR-A domains, a second map was generated following a different processing scheme, which used signal subtraction and is shown in Extended Data Fig. 2f. The final refinement led to a 3.8Å map of LRRK2$^{RCKW}$. All the steps used unbinned images (1.07Å $pixel^{-1}$).

The resolutions of the cryo-EM maps, here and below, were estimated from Fourier Shell Correlation (FSC) curves calculated using the gold-standard procedure and the resolutions are reported according to the 0.143 cutoff criterion[65,66]. FSC curves were corrected for the convolution effects of a soft mask applied to the half maps by high-resolution phase randomization[67]. For display and analysis purposes, we sharpened the maps with automatically estimated negative B factors from Relion or CryoSparc2.

We built the LRRK2$^{RCKW}$ models using both the 3.47Å C3 and 3.8Å density-subtracted maps. We used a combination of Rosetta[22] and manual building in Coot[68] to build all models. Starting models were found via a sequence alignment search in HHpred[69] and the top 5 results for each domain were used in the initial fitting of the backbone. We built the COR-B, KIN, and WD40 domains using the density of the 3.47Å C3 map and the RoC and COR-A domains using the 3.8Å map, then connected the two after fitting them into the 3.8Å map. Finally, we performed multiple iterations of both the CM and Relax functions in Rosetta, along with manual manipulation in Coot, to build our final 20 models (10 including GDP-$Mg^{2+}$ in the RoC domain, and 10 excluding it). In areas of weak density, we either removed part of the polypeptide chain or, where only side chain density was poor, converted the chain to poly-alanine.

The GDP-$Mg^{2+}$ was placed in our model by initially aligning a structure of the RoC domain containing a bound GDP-$Mg^{2+}$ (PDB: 2zej)[17] to the RoC domain in our structure. The GDP-$Mg^{2+}$ was then added to our model in the aligned position and run through Rosetta to allow for fine movements into our density and re-arrangement of nearby chains.

The phosphorylation of the threonine residue 1343, which we observe in our map and is a known phosphorylation site of Roco family GTPases[70], was confirmed by phosphor-enrichment mass spec (data not shown).

*Electron microscopy sample preparation, imaging, and processing of apo, MLi2, Ponatinib LRRK2$^{RCKW}$ monomer/dimers:*

For all samples, purified LRRK2$^{RCKW}$ was dialyzed into the same final buffer as described for the trimer data, then diluted to its final concentration in the same buffer. Unlike with the trimer data, however, the following datasets were collected from multiple grids prepared using slightly different sample conditions and imaged using a range of microscope settings.

The apo LRRK2$^{RCKW}$ monomer dataset had sample diluted to final concentrations ranging between 1 µM and 6 µM. In addition, one dataset was collected with the grid tilted to 30° to overcome preferred orientation issues. Otherwise, grids were prepared as described for the trimer dataset.

The apo LRRK2$^{RCKW}$ dimer dataset had sample diluted to final concentrations ranging between 4 µM and 12 µM. Two of the samples contained 0.05mM Digitonin (Sigma, D141) or 0.03% octyl glucoside (Sigma, O8001) detergents to overcome preferred orientation issues. One dataset was collected with the grid tilted to 30°, also to overcome preferred orientation issues. Otherwise, grids were prepared as described for the trimer dataset.

The MLi-2 LRRK2$^{RCKW}$ dimer dataset had sample diluted to final concentrations of either 3 µM or 4 µM. MLi-2 was added post-dialysis to a final concentration of 5 µM. The sample was incubated on ice for at least one hour before being applied to the grid. Otherwise, grids were prepared as described for the trimer dataset.

The ponatinib LRRK2$^{RCKW}$ dimer dataset had sample diluted to final concentrations of 2 μM or 4 μM. Ponatinib was added post-dialysis at a concentration of either 5 μM or 100 μM. The sample was incubated on ice for at least one hour before being applied to the grid. Otherwise, grids were prepared as described for the trimer dataset.

The apo LRRK2$^{RCKW}$ monomer cryo-EM data was collected on a Talos Arctica (FEI) operated at 200kV, equipped with a K2 Summit direct electron detector (Gatan). Automated data collection was performed using Leginon[58]. A total of 11,354 movies were collected. We imaged samples in both 'counting mode' and 'super resolution mode' at dose rates between 4.2 and 10 electrons $Å^{-2}$ $sec^{-1}$ with total exposure times ranging from 6s to 12s sub-divided into 200ms frames, for a total of 30 or 60 frames. All the images were recorded at nominal magnifications of either 36,000x (counting mode) or 62,000x (super resolution mode) resulting in object pixel sizes of either 1.16Å or 0.58Å, respectively. The defocus range of the data was -1 μm to -2 μm.

Frame alignment, CTF estimation and image selection were performed as described for the trimer dataset except that per-particle CTF was not used, instead the CTF information of the whole image was used. After selection we had 7,067 micrographs. Particles were extracted using crYOLO[71]. We carried out subsequent processing in CryoSparc2[64] on binned images (2.32Å $pixel^{-1}$).

A series of 2D and 3D classifications were performed as shown in Extended Data Fig. 8 to generate the final map. The initial monomer reference, used for refinement of the monomer map, was generated from our LRRK2$^{RCKW}$ model. The initial dimer references, used for particle sorting, were generated as shown in Extended Data Fig. 7. All references were filtered to 30Å, the default value in CrypSparc2, before refinement processes. This final map reached a resolution of 8.08Å using non-uniform refinement.

The apo LRRK2$^{RCKW}$ dimer cryo-EM data was collected as described for the apo LRRK2$^{RCKW}$ monomer. A total of 5,303 movies were collected. We imaged samples in both 'counting mode' and 'super resolution mode' at dose rates between 4.6 and 7.8 electrons $Å^{-2}$ $sec^{-1}$ with total exposure times ranging between 7s and 11s sub-divided into 200ms frames, for a total of 35 or 55 frames. All the images were recorded at nominal magnifications of either 36,000x (counting mode) or 62,000x (super resolution mode) resulting in object pixel sizes of either 1.16Å or 0.58Å, respectively. The defocus range of the data was -1 μm to -2 μm.

Frame alignment, CTF estimation, image selection, and particle picking were performed as described for the apo LRRK2$^{RCKW}$ monomer. After selection we had 3,100 micrographs. We carried out subsequent processing in CryoSparc2[64] on binned images (2.32Å $pixel^{-1}$).

The classification and refinement scheme for the apo LRRK2$^{RCKW}$ WD40- and COR-mediated dimer maps is shown in Extended Data Fig. 8. The same initial dimer references used for the apo LRRK2$^{RCKW}$ monomer, filtered to the same resolution, were used here for refinement of the dimer maps. In addition, a linear trimer reference, generated from the two initial dimer references and filtered to the same resolution, was used during the initial 3D classification step to sort out bad particles. The final maps had resolutions of 13.39Å (WD40-mediated dimer) and 9.52Å (COR-mediated dimer) with C2 symmetry applied to both.

The MLi-2 LRRK2$^{RCKW}$ dimer cryo-EM data was collected as described for the apo LRRK2$^{RCKW}$ monomer. We recorded a total of 4,139 movies. We imaged all datasets in 'counting mode' at a dose rate of 5.5 electrons $Å^{-2}$ $sec^{-1}$, with total exposure times of either 9s or 10s sub-divided into 200 ms frames, for a total of 45 or 50 frames. All the images were recorded at a nominal magnification of 36,000x (counting mode) resulting in object pixel sizes of 1.16Å. The defocus range of the data was -1 μm to -2 μm.

Frame alignment, CTF estimation, image selection, and particle picking were performed as described for the apo LRRK2$^{RCKW}$ monomer. After selection, 4,030 micrographs were kept for further processing. Processing was done in Relion 3.0[63] then CryoSparc2[64] using binned images (2.32Å $pixel^{-1}$).

The classification and refinement scheme for the MLi-2 LRRK2$^{RCKW}$ WD40- and COR-mediated dimer maps is shown in Extended Data Fig. 9. The same references used for the apo LRRK2$^{RCKW}$ dimers, and filtered to the same resolution, were used here for the same purposes. The final maps had resolutions of 9.74Å (WD40-mediated dimer) and 9.04Å (COR-mediated dimer), with no symmetry applied.

The Ponatinib LRRK2$^{RCKW}$ dimer cryo-EM data was collected as described for the apo LRRK2$^{RCKW}$ monomer. We recorded a total of 1,797 movies. We imaged all datasets in 'counting mode' at dose rates of either 5.5 or 9.7 electrons $Å^{-2}$ $sec^{-1}$ with total exposure times of 7s or 10s sub-divided into 200ms frames, for a total of 35 or 50 frames. All the images were recorded at a nominal magnification of 36,000x (counting mode) resulting in object pixel sizes of 1.16Å. The defocus range of the data was -1um to -2um.

Frame alignment, CTF estimation, image selection, and particle picking were performed as described for the apo LRRK2$^{RCKW}$ monomer. 1,455 micrographs were kept for further processing. Processing was done in CryoSparc2[64] on binned images (2.32Å $pixel^{-1}$). Ponatinib LRRK2$^{RCKW}$ dimer particles were sorted via 3D and 2D classification leading to the final 2D averages shown in Extended Data Fig. 10.

*Electron microscopy sample preparation and imaging of microtubule-bound LRRK2$^{RCKW}$:*

Purified LRRK2$^{RCKW}$ was dialyzed into the same buffer used for the trimer dataset with the addition of 20 μM Taxol and then diluted to a final concentration of 3 μM in the same buffer. Microtubules, made as previously described[57], were then added to a final concentration of 3 μM (tubulin dimer concentration). The mixture was incubated at room temperature for a minimum of 5 minutes before being applied to the grid. Grids were prepared as described for the trimer dataset except that Quantifoil C-flat 1.2/1.3 carbon open hole grids were used.

Cryo-EM data was collected on a Talos Arctica (FEI) operated at 200kV, equipped with a K2 Summit direct electron detector (Gatan) using Leginon [58]. We recorded a total of 10 movies. We imaged in 'super resolution mode' at a dose rate of 7.9 electrons $Å^{-2}$ $sec^{-1}$ with a total exposure time of 7s sub-divided into 200ms frames, for a total of 45 frames. All the images were recorded at a nominal magnification of 62,000x resulting in object pixel sizes of 0.58Å. The defocus range of the data was -1 μm to -2 μm. Images were aligned using MotionCor2.

*Building the molecular model of microtubule-associated LRRK2$^{RCKW}$ filaments*

Given that the WD40 densities are clearly identifiable in the sub-tomogram average of microtubule (MT)-associated LRRK2, we used these as a starting point for docking the structure of LRRK2$^{RCKW}$ into the cryo-ET map. First, a synthetic dimer of the WD40 domains from the LRRK2$^{RCKW}$ structure was generated by aligning them to a crystal structure of the isolated WD40 domain, which formed a dimer in the crystal (PDB: 6DLP) [18]. This synthetic dimer was then docked into the sub-tomogram average in Chimera, using the Fit in Map function with the options of filtering the structure to the resolution of the map (14Å) and optimizing correlation. A WD40 dimer was placed into each of the two corresponding densities present in the map. Then, four copies of the LRRK2$^{RCKW}$ structure were added by aligning their WD40 domains to those previously docked into the sub-tomogram average. The same procedure was followed to build the filament using the "closed" kinase model of LRRK2$^{RCKW}$ (see section below for how that model was generated).

Backbone clashes at the COR-mediated interface in the model filament were measured in Chimera (with default settings) after converting the four LRRK2$^{RCKW}$ monomers to poly-alanine models.

<u>Modeling a "closed" kinase version of LRRK2$^{RCKW}$</u>

In order to identify a good reference to model the closed state of LRRK2$^{RCKW}$'s kinase, we ran separate structural searches (using the DALI server) with the N- and C-lobes of LRRK2$^{RCKW}$'s kinase domain. We looked through the matches for a kinase that scored highly with both lobes, and whose structure is in a closed state. We selected Interleukin-2 inducible T-cell kinase (Itk) bound to an inhibitor as our reference (PDB: 3QGY) [54].

LRRK2$^{RCKW}$ was split at the junction between the N- and C-lobes of its kinase domain (L1949-A1950), resulting in one half containing the RoC, COR, and kinase (N-lobe) domains and another containing the kinase (C-lobe) and WD40 domains. The C-lobe of 3QGY was then aligned (in Chimera) to the C-lobe of LRRK2$^{RCKW}$'s kinase domain, and subsequently the N-lobe of LRRK2$^{RCKW}$'s kinase was aligned to the N-lobe of 3QGY. The two halves were then combined to generate the "closed" kinase model of LRRK2$^{RCKW}$.

<u>Docking of LRRK2$^{RCKW}$ into cryo-EM maps of monomers and dimers</u>

In order to build models of WD40- and COR-mediated dimers of LRRK2$^{RCKW}$ in the presence of MLi-2, we again split LRRK2$^{RCKW}$ at the junction between the N- and C-lobes (L1949-A1950). The two halves were fitted into one half of the cryo-EM map of a WD40-mediated dimer of LRRK2$^{RCKW}$ obtained in the presence of MLi-2 (we chose this map as its resolution was higher than that of the COR-mediated dimer). We also docked the two halves of LRRK2$^{RCKW}$ into a cryo-EM map of a LRRK2$^{RCKW}$ monomer obtained in the absence of inhibitor. The fitting was done in Chimera using the Fit in Map function with the options of filtering the structure to the resolution of the map and optimizing correlation. The two halves were then joined to generate a full model of LRRK2$^{RCKW}$.

The WD40- and COR-mediated dimers of LRRK2$^{RCKW}$ in the presence of MLi-2 were built by docking the models built above into the corresponding cryo-EM maps, using the same approach in Chimera as outlined above.

The LRRK2$^{RCKW}$ "filament" shown in Extended Data Fig. 11 was generated by aligning, in alternating order, multiple copies of the two dimer models (WD40- and COR-mediated) built into the cryo-EM maps obtained in the presence of MLi-2.

Kinase inhibitors

Stocks of the kinase inhibitors MLi-2 (10 mM; Tocris), Ponatinib (10 mM; ApexBio), GZD-824 (10 mM; Cayman Chemical), and LRRK2-IN-1 (2 mM; Michael J Fox Foundation) were stored in DMSO at -20°C.

Antibodies

All antibodies used for immunocytochemistry were diluted to 1:500. Primary antibodies used were chicken anti-GFP (Aves Labs) and rabbit anti-alpha-tubulin (ProteichTech). Secondary antibodies used were goat anti-chicken-Alexa 488 (ThermoFisher) and goat anti-chicken Alexa568 (ThermoFisher). DAPI was used at 1:5000 according to the manufacturers suggestions (ThermoFisher). Primary antibodies used for Western blots were mouse anti-GFP (Santa Cruz, 1:1000 dilution) mouse anti-GAPDH (ProteinTech, 1:5000 dilution) and mouse anti-gamma-tubulin (ProteinTech, 1:5000 dilution). Secondary antibodies (1:15,000) used for Western blots were IRDye goat anti-mouse 680RD and IRDye goat anti-rabbit 780RD (Li-COR).

Rab8a expression and purification

N-terminally tagged ($His_6$-ZZ) Rab8a containing a TEV cleavage site was cloned into a PET28a expression vector and expressed in BL21(DE3) *E. coli* cells. Transformed cells were grown overnight at 37ºC in 10 mL LB medium containing kanamycin (50 µg/ml), then diluted into 200 mL LB medium containing kanamycin (50 µg/ml), grown to an optical density at 600 nm of ~1-2, diluted into 4 L LB medium containing kanamycin (50 µg/ml), and grown to an optical density at 600 nm of 0.4. IPTG was added (final concentration 0.5 mM) to induce protein expression for ~18 hours at 18ºC. Cells were harvested by centrifugation at 8983 x g for 10 min at 4ºC, followed by resuspension in 15 mL LB medium and centrifugation at 2862 x g for 10 min at 4ºC. The cell pellet was flash frozen in liquid nitrogen and stored at -80ºC. For a typical protein purification, cell pellets were resuspended in lysis buffer (50 mM HEPES pH 7.4, 200 mM NaCl, 2 mM DTT, 10% glycerol, 5 mM $MgCl_2$, 0.5 mM Pefabloc, and protease inhibitor cocktail tablets) and lysed by sonication on ice. The lysate was clarified by centrifugation at 164,700 x g for 40 mins at 4ºC and then incubated with Ni-NTA agarose beads (Qiagen) for 1 hour at 4ºC. Beads were extensively washed with wash buffer (50 mM HEPES pH 7.4, 150 mM NaCl, 2 mM DTT, 10% glycerol, 5 mM $MgCl_2$); $His_6$-ZZ-Rab8a was eluted in 40 mL elution buffer (50 mM HEPES pH 7.4, 150 mM NaCl, 300 mM imidazole, 2 mM DTT, 10% glycerol, 5 mM $MgCl_2$). The protein eluate was diluted 2 fold in wash buffer, incubated with IgG sepharose 6 fast flow beads equilibrated in wash buffer, incubated at 4ºC for 2.5 hours, and washed extensively in wash buffer. Protein-bound IgG beads were then transferred into TEV buffer (50 mM HEPES pH 7.4, 200 mM NaCl, 2 mM DTT, 10% glycerol, 5 mM $MgCl_2$), and untagged Rab8a was cleaved off of IgG sepharose beads by incubation with TEV protease at 4ºC overnight. The next day, cleaved Rab8a was separated from $His_6$-TEV protease and any remaining uncleaved protein or residual tag by incubation with Ni-NTA agarose beads (Qiagen), followed by washing with TEV buffer containing 25 mM imidazole. Lastly, purified Rab8a was run over a Superdex 200 increase 10/300 size exclusion column equilibrated in S200 buffer (50 mM HEPES pH 7.4, 200 mM NaCl, 2 mM DTT, 1% glycerol, 5 mM $MgCl_2$), and concentrated and exchanged into buffer containing 10% glycerol for storage at -80ºC.

*In vitro* phosphorylation of Rab8a by LRRK2$^{RCKW}$

Purified Rab8a (~3.8 µM) was phosphorylated by LRRK2$^{RCKW}$ (~38 nM) in a buffer containing 50 mM HEPES pH 7.4, 80 mM NaCl, 10 mM $MgCl_2$, 0.5 mM TCEP, 1 mM ATP, 200 µM GDP); 34 µL reaction mixtures containing kinase inhibitor or an equivalent volume DMSO were incubated at 30ºC, and samples were taken at 45 mins, and 90 mins. An effective reaction volume of 0.75 µL was run on a 4-12% Bis-Tris protein gel, transferred to nitrocellulose, and blotted with a commercially available antibody to pT72-Rab8a (MJFF-pRab8) as previously described[72] and per manufacturer's instructions, with the exception that HRP-labeled secondary antibody was used at a dilution of 1:2000.

Purification of molecular motors

*Kinesin*

Protein purification steps were done at 4°C unless otherwise indicated. Human KIF5B$^{1\text{-}560}$ (K560)-GFP was purified from *E. coli* using an adapted protocol previously described[73]. pET17b-Kif5b(1-560)-GFP-His was transformed into BL-21[DE3] RIPL cells (New England Biolabs) until OD 0.6-0.8 and expression was induced with 0.5 mM IPTG for 16 hr at 18°C. Frozen pellets from 2 L culture were resuspended in 40 mL lysis buffer (50 mM Tris, 300 mM NaCl, 5 mM $MgCl_2$, and 0.2 M sucrose, pH 7.5) supplemented with 1 cOmplete EDTA-free protease inhibitor cocktail tablet (Roche) per 50 mL and 1 mg/mL lysozyme. The resuspension was incubated on ice for 30 min and lysed by sonication. Sonicate was supplied with 10 mM imidazole and 0.5 mM PMSF and clarified by centrifuging at 30,000 x g for 30 min in Type 70 Ti rotor (Beckman). The clarified supernatant was incubated with 5 mL Ni-NTA agarose (Qiagen) and rotated in a nutator for 1 hr. The mixture was washed with 30 mL wash buffer (50 mM Tris, 300 mM NaCl, 5 mM $MgCl_2$, 0.2 M sucrose, and 20 mM imidazole, pH 7.5.) by gravity flow. Beads were resuspended in elution buffer (50 mM Tris, 300 mM NaCl, 5 mM $MgCl_2$, 0.2 M sucrose, and 250 mM imidazole, pH 8.0), incubated for 5 mins, and eluted stepwise in 0.5 mL increments. Peak fractions were combined and buffer exchanged on a PD-10 desalting column (GE Healthcare) equilibrated with storage buffer (80 mM PIPES, 2 mM $MgCl_2$, 1 mM EGTA, and 0.2 M sucrose, pH 7.0). From this, peak fractions of motor solution were either flash frozen at -80°C until further use or immediately subjected to microtubule bind and release purification. A total of 1 mL motor solution was incubated with 1 mM AMP-PNP and 20 µM taxol on ice for 5 mins and warmed to room temperature (RT). For microtubule bind and release, polymerized bovine brain tubulin was centrifuged through a glycerol cushion (80 mM PIPES, 2 mM $MgCl_2$, 1 mM EGTA, and 60 % glycerol (v/v) with 20 µM taxol and 1 mM DTT) and resuspended as previously described[32] was incubated with motor solution in the dark for 15 mins at RT. The Motor-microtubule mixture was laid on top of a glycerol centrifuged in a TLA120.2 rotor at 278,835 x g for 12 min at RT. Final pellet (Kinesin-bound microtubules) was washed with BRB80 (80 mM PIPES, 2 mM $MgCl_2$, and 1 mM EGTA, pH 7.0) and incubated in 100 µL of release buffer (80 mM PIPES, 2 mM $MgCl_2$, 1 mM EGTA, and 300 mM KCl, pH ~7 with 5mM Mg-ATP) for 5 mins at RT. The supernatant was supplied with 660 mM sucrose and flash frozen. A typical kinesin prep yielded ~0.5 to 1 µM K560-GFP dimer.

*Dynein*

Human dynactin was purified from stable cell lines expressing p62-Halo-3xFlag as described previously[74]. Briefly, cells were collected from 160 x 15 cm plates and resuspended in 80 mL of dynactin-lysis buffer (30 mM HEPES [pH 7.4], 50 mM potassium acetate, 2 mM magnesium acetate, 1 mM EGTA, 1 mM DTT, 10% (v/v) glycerol) supplemented with 0.5 mM Mg-ATP, 0.2% Triton X-100 and 1 cOmplete EDTA-free protease inhibitor cocktail tablet (Roche) per 50 mL and rotated slowly for 15 min. The lysate was clarified by centrifuging at 66,000 x g for 30 min in Type 70 Ti rotor (Beckman). The clarified supernatant was incubated with 1.5 mL of anti-Flag M2 affinity gel (Sigma-Aldrich) overnight on a roller. The beads were transferred to a gravity flow column, washed with 50 mL of wash buffer (dynactin-lysis buffer supplemented with 0.1 mM Mg-ATP, 0.5 mM Pefabloc and 0.02% Triton X-100), 100 mL of wash buffer supplemented with 250 mM potassium acetate, and again with 100 mL of wash buffer. Dynactin was eluted from beads with 1 mL of elution buffer (wash buffer with 2 mg/mL of 3xFlag peptide). The eluate was collected, filtered by centrifuging with Ultrafree-MC VV filter (EMD Millipore) in a tabletop centrifuge and diluted to 2 mL in Buffer A (50 mM Tris-HCl [pH 8.0], 2 mM MgOAc, 1 mM EGTA, and 1 mM DTT) and injected onto a MonoQ 5/50 GL column (GE Healthcare and Life Sciences) at 1 mL/min. The column was pre-washed with 10 column volumes (CV) of Buffer A, 10 CV of Buffer B (50 mM Tris-HCl [pH 8.0], 2 mM MgOAc, 1 mM EGTA, 1 mM DTT, 1 M KOAc) and again with 10 CV of Buffer A at 1 mL/min. To elute, a linear gradient was run over 26 CV from 35-100% Buffer B. Pure dynactin complex eluted from ~75-80% Buffer B. Peak fractions containing pure dynactin complex were pooled, buffer exchanged into a GF150 buffer supplemented with 10% glycerol, concentrated to 0.02-0.1 mg/mL using a 100K MWCO concentrator (EMD Millipore) and flash frozen in liquid nitrogen. Typical dynactin prep yields are between 150-300 nM.

Human dynein was purified from stable cells lines expressing an IC2-SNAPf-3xFlag as described previously[33]. Frozen pellets collected from ~60-100 x 15 cm plates were resuspended in dynein lysis buffer (25 mM HEPES pH 7.4, 50 mM KOAc, 2 mM MgOAc,1 mM EGTA, 10% glycerol (v/v), and 1 mM DTT) supplemented with 0.2% Triton X-100, 0.5 mM Mg-ATP, and cOmplete EDTA-free protease inhibitor cocktail. The lysate was centrifuged at 66,000 x g in a Ti-70 rotor for 30 mins. The clarified supernatant was incubated with 1 mL of anti-Flag M2 affinity gel (Sigma-Aldrich) overnight on a roller. Beads were collected by gravity flow and washed with 50 mL wash buffer (dynein lysis buffer with 0.02% Triton X-100 and 0.5 mM Mg-ATP) supplemented with protease inhibitors (cOmplete Protease Inhibitor Cocktail, Roche). Beads were then washed with 50 mL high salt wash buffer (25 mM HEPES, pH 7.4, 300 mM KOAc, 2 mM MgOAc, 10% glycerol, 1 mM DTT, 0.02% Triton X-100, 0.5 mM Mg-ATP), and then with 100 mL wash buffer. For labeling, beads were resuspended in 1 mL wash buffer and incubated with 5 µM SNAP-Cell TMR Star (New England BioLabs) for 10 min on the column at RT. Unbound dye was removed with 100 mL wash buffer at 4°C. Dynein was eluted with 1 mL of elution buffer (wash buffer containing 2 mg/mL 3xFLAG peptide). The eluate was collected, diluted to 2 mL in Buffer A (50 mM Tris pH 8.0, 2 mM MgOAc, 1 mM EGTA, and 1 mM DTT) and injected onto a MonoQ 5/50 GL column (GE Healthcare Life Sciences) at 0.5 mL/min. The column was washed with 20 CV of Buffer A at 1 mL/min. To elute, a linear gradient was run over 40 CV into Buffer B (50 mM Tris pH 8.0, 2 mM MgOAc, 1 mM EGTA, 1 mM DTT, 1 M KOAc). Pure dynein complex elutes from ~60–70% Buffer B. Peak fractions were pooled and concentrated, 0.1 mM Mg-ATP and 10% glycerol were added and the samples were snap frozen in liquid nitrogen. A typical preparation yielded 150-300 nM dynein.

*NINL*

Human NINL was purified as previously described[74]. pET28a-ZZ-TEV-Halo-NINL$^{1-702}$ was transformed into BL-21[DE3] cells (New England Biolabs) until OD 0.4-0.6 and expression was induced with 0.1 mM IPTG for 16 hr at 18°C. Frozen cell pellets from 1 L culture were resuspended in 40 mL of activator-lysis buffer (30 mM HEPES [pH 7.4], 50 mM potassium acetate, 2 mM magnesium acetate, 1 mM EGTA, 1 mM DTT, 0.5 mM Pefabloc, 10% (v/v) glycerol) supplemented with 1 cOmplete EDTA-free protease inhibitor cocktail tablet (Roche) per 50 mL and 1 mg/mL lysozyme. The resuspension was incubated on ice for 30 min and lysed by sonication. The lysate was clarified by centrifuging at 66,000 x g for 30 min in Type 70 Ti rotor (Beckman). The clarified supernatant was incubated with 2 mL of packed IgG Sepharose 6 Fast Flow beads (GE Healthcare Life Sciences) for 2 hr on a roller. The beads were transferred to a gravity flow column, washed with 100 mL of activator-lysis buffer supplemented with 150 mM potassium acetate and 50 mL of cleavage buffer (50 mM Tris–HCl [pH 8.0], 150 mM potassium acetate, 2 mM magnesium acetate, 1 mM EGTA, 1 mM DTT, 0.5 mM Pefabloc, 10% (v/v) glycerol). The beads were then resuspended and incubated in 15 mL of cleavage buffer supplemented with 0.2 mg/mL TEV protease overnight on a roller. The supernatant containing cleaved proteins were concentrated using a 50K MWCO concentrator (EMD Millipore) to 1 mL, filtered by centrifuging with Ultrafree-MC VV filter (EMD Millipore) in a tabletop centrifuge, diluted to 2 mL in Buffer A (30 mM HEPES [pH 7.4], 50 mM potassium acetate, 2 mM magnesium acetate, 1 mM EGTA, 10% (v/v) glycerol and 1 mM DTT) and injected onto a MonoQ 5/50 GL column (GE Healthcare and Life Sciences) at 0.5 mL/min. The column was pre-washed with 10 CV of Buffer A, 10 CV of Buffer B (30 mM HEPES [pH 7.4], 1 M potassium acetate, 2 mM magnesium acetate, 1 mM EGTA, 10% (v/v) glycerol and 1 mM DTT) and again with 10 CV of Buffer A at 1 mL/min. To elute, a linear gradient was run over 26 CV from 0-100% Buffer B. The peak fractions containing unlabeled Halo-tagged NINL were collected and concentrated to using a 50K MWCO concentrator (EMD Millipore) to 0.2 mL. A typical NINL prep yield was ~5-10 µM dimer.

Single-molecule microscopy and motility assays

Single-molecule imaging was performed using total internal reflection fluorescence (TIRF) microscopy with an inverted microscope (Nikon, Ti-E Eclipse) equipped with a 100x 1.49 N.A. oil immersion objective (Nikon, Plano Apo), and a MLC400B laser launch (Agilent), with 405 nm, 488 nm, 561 nm and 640 nm laser lines. Excitation and emission paths were filtered using single bandpass filter cubes (Chroma), and emitted signals were detected with an electron multiplying CCD camera (Andor Technology, iXon Ultra 888). Illumination and image acquisition were controlled with NIS Elements Advanced Research software (Nikon), and the xy position of the stage was controlled with a ProScan linear motor stage controller (Prior).

Single-molecule motility were performed in flow chambers as previously described[33] using the setup shown in the schematic in Fig. 4A. Biotin-PEG-functionalized coverslips (Microsurfaces) were adhered to a Superfrost Plus Microscope slide (ThermoFisher) using double-sided scotch tape. Each slide contained four flow-chambers. Taxol-stabilized microtubules (~15mg/mL) with ~10% biotin-tubulin and ~10% Alex405-tubulin were prepared as described previously[33]. For each motility experiment, 1 mg/mL Strepdavidin (in 30 mM HEPES, 2 mM MgOAc, 1mM EGTA, 10% glycerol) was incubated in the flow chamber for 3 mins. A 1:150 dilution of taxol-stabilized microtubules in motility assay buffer (30 mM HEPES, 50 mM KOAc, 2 mM MgOAc, 1mM EGTA, 10% glycerol, 1 mM DTT, and 20 µM Taxol, pH 7.4) was added to the flow chamber for 3 mins to adhere polymerized microtubules to the coverslip. Flow chambers containing adhered microtubules were washed twice with LRRK2 buffer (20 mM HEPES pH 7.4, 800 mM NaCl, 0.5 mM TCEP, 5% glycerol, 2.5 mM $MgCl_2$, 20 µM GDP). Flow chambers were then incubated for 5 mins either with (1) LRRK2 buffer alone or with the indicated kinase inhibitors ("0 nM" LRRK2$^{RCKW}$ condition) or (2) LRRK2 buffer containing LRRK2$^{RCKW}$ either alone, with DMSO, or with kinase inhibitors. DMSO or drugs were incubated with LRRK2 buffer (± LRRK2$^{RCKW}$) for 10 mins at RT before adding to the flow chambers. Prior to the addition of dynein and kinesin motors, the flow chambers were washed twice with motility assay buffer containing 1mg/mL casein. To assemble dynein-dynactin-ninein-like (NINL) complexes, purified dynein (10-15 nM), dynactin and NINL were mixed at 1:2:10 molar ratio and incubated on ice for 10 min. The final imaging buffer for motors contained motility assay buffer supplemented with an oxygen scavenger system, 71.5 mM β-mercaptoethanol and either 1 mM ATP (kinesin) or 2.5 mM ATP (dynein). The final concentrations of kinesin and dynein in the flow chambers were ~2.5 nM and ~0.3 nM, respectively. K560-GFP was imaged every 500 msecs for 2 mins with 25% laser (488) power at 150 ms exposure time. Dynein-TMR-dynactin-NINL was imaged every 300 msec for 3 mins with 25% laser (561) power at 100 msec. Each sample was imaged no longer than 15 mins. Each technical replicate consisted of movies from at least two fields of view containing between 5 and 10 microtubules each.

Single-molecule motility assay analysis

Kymographs were generated from motility movies and quantified for run lengths, percent motility, and velocity using ImageJ (NIH). Specifically, maximum-intensity projections were generated from time-lapse sequences to define the trajectory of particles on a single microtubule. The segmented line tool was used to trace the trajectories and map them onto the original video sequence, which was subsequently re-sliced to generate a kymograph. Motile and immotile events (> 1 sec) were manually traced. Bright aggregates, which were less than 5% of the population, were excluded from the analysis. For dynein-dynactin-NINL, both stationary and diffusive events were grouped as immotile. Run length measurements were calculated from motile events only. For percent motility per microtubule measurements, motile events (> 1 sec and > 1µm) were divided by total events per kymograph. Velocity measurements were calculated from the inverse slopes of the motile event traces (> 1 sec and > 1µm) only. Statistical analyses were performed in Prism8 (Graphpad).

Western blot analysis

293T cells were maintained in Dulbecco's modified Eagle's medium (containing 10% fetal bovine serum and 1% penicillin/streptomycin). For Western blot quantification of LRRK2 protein expression (Extended Data Fig. 15), cells were plated on 6-well dishes (150K cells per well) 24 hrs before transfection. Cells were transfected with 1 µg of GFP-I2020T using polyethylenimine (PEI, Polysciences). After 48 hrs, cells were treated for 30 mins with either 5 µM GZD-824 or DMSO-matched control. Cells were lysed on ice in RIPA buffer (50 mM Tris pH 7.5, 150 mM NaCl, 0.2% TritonX-10, 0.1%SDS, 0.5% Na-Deoxycholate, with cOmplete protease inhibitor cocktail). Lysates were further rotated for 15 mins at 4°C and clarified by centrifuging at 13,000 x g for 15 mins. Clarified supernatants were boiled for 5 mins in Laemmli buffer. The experiments were performed in triplicate.

For Western blots, lysates were run on 4-12% gradient SDS-PAGE (Life Technologies) for 60 mins and transferred to nitrocellulose for 3 hrs at 250 mA. Blots were dried at RT for 30 mins, rinsed in 1x Tris buffered saline (TBS), followed by blocking with 5% milk in TBS. Antibodies were diluted in 5% milk in TBS-0.1% Tween-20 (TBS-T). Primary antibodies were incubated overnight at 4°C and Infrared (IR) secondary antibodies were incubated at RT for 45 mins. For quantification of LRRK2 expression levels, blots were imaged on an Odyssey CLx controlled by Imaging Studio software (v5.2). DMSO and GZD-824 conditions were quantified in triplicate and normalized to a GAPDH loading control using Empiria Studio software (Li-COR). To ensure quantification was in the combined linear range for antibodies detecting both GFP-LRRK2 and GAPDH, a linear dilution series of lysates from cells expressing GFP-LRRK2 was also quantified by IR Western blot.

Immunofluorescence, confocal microscopy and image analysis

The day before transfection, 293T cells were plated on acid-treated coverslips (Bellco Glass) pre-coated with 100 µg/mL Poly-D-lysine (Sigma) and 4 µg/mL Mouse Laminin (ThermoFisher) in 24-well plates (35K cells per well). Cells were transfected with 500 ng plasmid of either pDEST53-GFP-LRRK2 or pDEST53-GFP-LRRK2(I2020T) using PEI. After 48-72 hrs, cells were incubated with either a kinase inhibitor or DMSO-matched control (matched for time and concentration). For the Type 1 inhibitor experiment, cells were incubated with DMSO or MLi-2 (500 nM) for 2 hrs. For the Type 2 inhibitor, cells were incubated with DMSO or GZD-824 (5 or 10 µM) for 30 mins. Cells were quickly washed 1x on ice with ice-cold PBS, and fixed with ice-cold 4%PFA/90%Methanol/5mM sodium bicarbonate for 10 mins at -20°C. Following fixation, the wells were immediately washed 3x with ice-cold PBS. Blocking buffer (1% BSA, 5% normal goat serum, 0.3%TritonX-100 in PBS) was added for 1 hr at RT. Primary antibodies were diluted (1:500) in antibody dilution buffer (1% BSA, 0.1% TritonX-100 in PBS) and incubated overnight at 4°C. After overnight incubation, the wells were washed 3x in PBS and incubated with secondary antibodies (1:500) in antibody dilution buffer for 1 hr at RT. After secondary incubation, the wells were washed 3x in PBS, 1x in ddH2O and mounted using CitiFluor AF2

(EMS) on Superfrost Plus Microscope slides (ThermoFisher). Coverslips were sealed with nail polish and stored at 4°C.

For the LRRK2 filament analysis in Fig. 5, experimenters were blinded to condition for both the imaging acquisition and analysis. Cells were imaged using a Nikon A1R HD confocal microscope with a LUN-V laser engine (405nm, 488nm, 561nm, and 640nm) and DU4 detector using bandpass and longpass filters for each channel (450/50, 525/50, 595/50, and 700/75). Slides were imaged on a Nikon Ti2 body using an Apo 60x 1.49 NA objective. Image stacks were acquired in resonant scanning mode with bidirectional scanning and 4x line averaging and 1.2 airy units. The lasers used were 405 nm, 488 nm and 561 nm. Illumination and image acquisition were controlled by NIS Elements Advanced Research software (Nikon Instruments). ImageJ was used to quantify the percent of cells with LRRK2 filaments. Maximum-intensity projections were generated from *z*-stack confocal images. Using the GFP immunofluorescence signal, transfected cells were traced. Cells were scored for the presence or absence of filaments using both the z-projection and z-stack micrographs as a guide. The presence of filaments was scored if the cells had either (a) a GFP filament signal greater than 5 µm or (b) bundles of filaments with at least two identifiable crosses. To calculate the percent cells with filaments, the number of cells with filaments was divided by the total number of transfected cells per technical (defined as one 24-well coverslip). Approximately 20 cells were quantified per replicate for each condition in Figure 5D (DMSO v. MLi-2) and between 40-100 cells were quantified per replicate for each condition in Figure 5F (DMSO v. GZD-824). The quantification of all cellular experiments comes from data collected on three separate days except for the 10 µM GZD-824 condition in Figure 5F which was performed on two separate days. All statistical analyses were performed in Prism8 (Graphpad).

### Methods and Extended Data References


53. Sui, H. & Downing, K. H. Structural basis of interprotofilament interaction and lateral deformation of microtubules. *Structure* **18**, 1022–1031 (2010).

54. Charrier, J.-D., Miller, A., Kay, D. P., Brenchley, G., Twin, H. C., Collier, P. N., Ramaya, S., Keily, S. B., Durrant, S. J., Knegtel, R. M. A., Tanner, A. J., Brown, K., Curnock, A. P. & Jimenez, J.-M. Discovery and structure-activity relationship of 3-aminopyrid-2-ones as potent and selective interleukin-2 inducible T-cell kinase (Itk) inhibitors. *J. Med. Chem.* **54,** 2341–2350 (2011).

55. Pellegrini, E., Signor, L., Singh, S., Boeri Erba, E. & Cusack, S. Structures of the inactive and active states of RIP2 kinase inform on the mechanism of activation. *PLoS ONE* **12,** e0177161 (2017).


Data availability

Cryo-EM Maps and molecular models have been deposited in the wwPDB and will be available upon publication.
